# TRMT6/61A-mediated m^1^A methylation facilitates human pre-tRNA maturation and prevents surveillance by XRN2

**DOI:** 10.64898/2026.05.17.725798

**Authors:** Xisheng Liu, Nhi Yen Tran Nguyen, Abigail Grace Johnston, Ilias Skeparnias, Monima Anam, Katherine Marlow, Naira Rehman, Andrew H. Kellum, Yinsheng Wang, Jinwei Zhang, Zhangli Su

## Abstract

Transfer RNAs (tRNAs) are dynamically regulated by RNA modifications. The conserved TRMT6/61A catalyzes m^1^A (*N*1-methyladenosine) deposition at position 58. While TRMT6/61A dysregulation is linked to human diseases, its downstream processing consequences and molecular surveillance mechanisms remain unclear. Here we demonstrate that TRMT6/61A installs m^1^ A on precursor tRNAs prior to processing. Utilizing a dTAG rapid depletion system, we show that acute loss of TRMT6/61A swiftly reprograms the human tRNAome. Although elongator tRNA fluctuations are buffered by isodecoder redundancy, hypomethylated tRNA^iMet^ is selectively and rapidly degraded by the exoribonuclease XRN2, reducing global protein synthesis and activating ATF4 expression. Furthermore, m^1^A_58_ is a prerequisite for tRNA end processing; its absence leads to the aberrant accumulation of unprocessed pre-tRNAs and disrupted tRNA-derived fragment (tRF) populations. Mechanistically, TRMT6/61A facilitates *in vitro* RNase Z cleavage, likely by promoting proper pre-tRNA folding. Lastly, XRN2 inhibition rescues tRNA^iMet^ levels and reverses growth defects, identifying the XRN2-mediated surveillance of tRNA^iMet^ as a primary driver of the cellular pathology. Collectively, our results uncover a pivotal role for TRMT6/61A-dependent m^1^A in human tRNA maturation and define the molecular checkpoints essential for translational homeostasis.

## Introduction

tRNA is well established for its essential role in translation. We now know that the tRNAome (the cell’s complete set of tRNA molecules) is not static. Instead, it is actively regulated by chemical modifications, processing, fragmentation, and subcellular localization in response to nutrient availability, environmental stress and developmental cues^1^. This dynamic regulation is critical for human health, and its dysregulation is directly linked to a wide range of human diseases^2, 3^. tRNAs are heavily modified, estimated to have 13 modifications per molecule – these modifications are crucial for their decoding activity and stability^4, 5, 6, 7, 8, 9, 10, 11^. The dysregulation of tRNA-modifying factors is a growing area of research, now termed “tRNA modopathy”^12, 13, 14, 15, 16^. This dysregulation can result from loss-of-function mutations, often leading to neurodevelopmental or metabolic disorders, or from increased expression of the enzymes, a frequent observation in cancers. For example, over-expressing a single tRNA-modifying enzyme METTL1 alone is sufficient to drive oncogenesis by stabilizing specific arginine tRNAs; over-expressing the specific arginine tRNA substrate alone is also sufficient to drive oncogenesis^17, 18^, highlighting the importance of studying these factors’ specificity in human disease. Understanding how tRNA modifications are regulated and their molecular functions has profound implications in human health.

Among these crucial modifications, *N*1-methyladenosine (m^1^A) stands out due to its diverse RNA substrates, dynamic regulation, and functional implications^19^. m^1^A blocks base pairing and introduces a positive charge to adenosine^20, 21^. It is dynamically regulated in response to stress and different biological conditions^22, 23, 24, 25, 26, 27^. Accumulating evidence shows that m^1^A writers and erasers contribute to the dynamic regulation of m^1^A on different RNA substrates^25, 28, 29^. TRMT6/61A is the only known writer complex for catalyzing m^1^A at the 58^th^ position on tRNA T-arm, a modification conserved from yeast to human. TRMT6/61A is over-expressed in multiple cancer types and its inhibition is a promising therapeutic target^30, 31, 32, 33, 34, 35^. In addition, homozygous missense mutation in the catalytic subunit TRMT61A was reported in atypical De Lange Syndrome patient^36^.

Despite the conservation of m^1^A_58_, the molecular pathways governing its influence on the human tRNAome remain unclear. While studies in yeast highlight a selective destabilization of tRNA^iMet^ via the 5’-3’ rapid tRNA decay (RTD) and 3’-5’ nuclear surveillance pathways ^37, 38, 39, 40, 41^, mammalian responses to TRMT6/61A dysregulation appear context-dependent^31, 32, 33, 34, 35^. For instance, TRMT6/61A-mediated m^1^A on tRNA^Leu^ and tRNA^Ser^ has been shown to modulate *Myc* translation in T cells^34^; while in colon cancer cells, it regulates histone synthesis via the stability of tRNA-Lys-TTT-1^35^. Reconciling these observations is a significant challenge, as existing studies primarily rely on long-term genetic depletion (conditional knock-out or knock-down). Such chronic loss inevitably triggers secondary and compensatory mechanisms, growth defects, and stress signaling that obscure the primary, direct targets of the TRMT6/61A complex.

Furthermore, a critical but overlooked dimension of this regulation is the early stage of the tRNA lifecycle. Mature tRNAs are the product of an orchestrated maturation program, including 5’ leader removal by RNase P, 3’ trailer cleavage by RNase Z, 3’ CCA addition, and rigorous quality control. Yet it remains unknown whether TRMT6/61A acts upstream or downstream from these processing steps. To disentangle these immediate molecular events from long-term cellular adaptation, we engineered degradation TAG (dTAG) human cell lines to achieve rapid depletion of TRMT6/61A. By capturing the primary response of the tRNAome, we uncover a pivotal role for TRMT6/61A-mediated m^1^A in facilitating human pre-tRNA processing and protecting the initiator tRNA from rapid decay surveillance.

## Results

### Rapid degradation of endogenous TRMT6/61A by dTAG

To track temporal changes upon the loss of TRMT6/61A, we constructed dTAG-TRMT6 and dTAG-TRMT61A knock-in HEK293T cell lines via CRISPR/Cas9 homology-directed repair strategy (**Fig. 1a**). The dTAG cassette contains the mutant FKBP12^F36V^ tag to recruit the endogenous CRBN E3 ligase to the target protein upon addition of heterobifunctional degraders^42^. Successful insertion of the knock-in cassette at the N-terminus of the endogenous genes was confirmed by PCR using flanking primer pairs (**Fig. 1b**) and by Western blotting (**Fig. 1c-d**). Next, we tested different doses of dTAG-13 to identify the optimal concentration for rapid degradation of tagged TRMT6 within a 4-hour window (**Fig. 1e**). Both 0.1 and 0.5 µM dTAG-13 effectively induced degradation, while lower doses were insufficient. Interestingly, higher dTAG concentration did not improve degradation, which is consistent with the previously described “hook effect”^42^. To evaluate whether the dTAG insertion perturbs the functions of the TRMT6/61A complex, we confirmed the nuclear localization of tagged TRMT6 (**Fig. 1f**), as well as TRMT6/61A protein-protein interaction by co-immunoprecipitation (**Supplementary Fig. 1b**).

**Figure 1.**
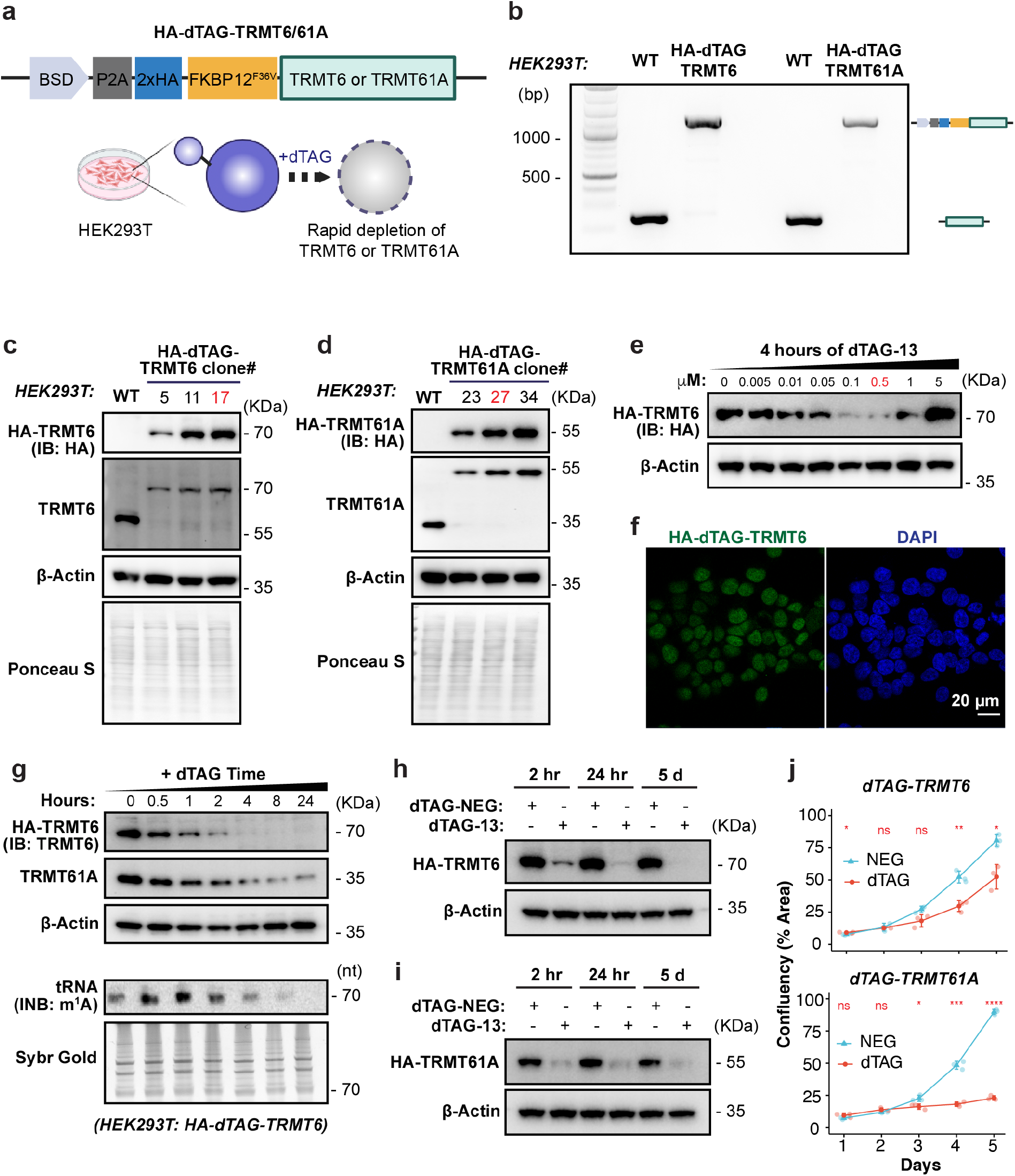
Rapid degradation of endogenous TRMT6/61A by dTAG. **(a)** Scheme of HA-dTAG-TRMT6/61A cell line construction. **(b)** Genomic PCR to confirm successful knock-in in HEK293T cells after blasticidin selection. **(c-d)** Western blotting to confirm successful knock-in in HEK293T individual clones. **(e)** Titration of dTAG-13 to determine the optimal concentration for protein degradation. **(f)** Immunofluorescence of HA-dTAG-TRMT6 in HEK293T. **(g-h)** Time course of 0.5 µM dTAG treatment in HA-dTAG-TRMT6 cells. **(i)** Time course of 0.5 µM dTAG treatment in HA-dTAG-TRMT61A cells. **(j)** Cell growth is reduced by dTAG-induced TRMT6/61A loss in HEK293T cells (n = 3). See also **Supplementary Figure 1**.

Using a fixed concentration of 0.5 µM dTAG-13, we followed the dynamics of TRMT6 degradation (**Fig. 1g-h**). To control for potential off-target and toxic effects from dTAG compound, we included the dTAG-13-NEG compound as a control in all subsequent experiments; this compound is a stereoisomer of dTAG-13 that cannot recruit the E3 ligase. Upon targeting TRMT6 with dTAG, endogenous TRMT6 protein levels gradually decreased to ∼10% by four hours. This was accompanied by a slower decrease in TRMT61A protein levels down to ∼20% by eight hours. A similar trend was observed when TRMT61A is targeted for degradation: as it was degraded to ∼10% by eight hours, its partner TRMT6 was also destabilized to ∼65% (**Fig. 1i** and **Supplementary Fig. 1c**). This suggests that the TRMT6/61A complex stabilizes both subunits, and that TRMT61A stability is more dependent on this interaction, implying that TRMT6 might have a longer half-life due to intrinsic protein properties or other interactors.

Interestingly, while tRNA m^1^A levels exhibited a gradual decrease following four hours of dTAG treatment, the initial phase (30 minutes to two hours) revealed a marked increase in m^1^A even as the writer was being degraded (**Fig. 1g**). This paradoxical observation might suggest an initial stress response. Eventually, the prolonged loss of TRMT6/61A for five days caused a significant decrease in cell proliferation (**Fig. 1j**). We also applied the N-terminal dTAG strategy to construct dTAG-TRMT6 and dTAG-TRMT61A HAP1 cell lines (**Supplementary Fig. 1d**), which displayed similarly rapid protein degradation in two hours (**Supplementary Fig. 1e-f**). Collectively, these dTAG models provide a robust, rapidly inducible system to dissect the temporal biological functions of the TRMT6/61A complex.

### Loss of TRMT6/61A causes rapid global tRNAome remodeling

Based on the dTAG degradation timeline, we selected three time points to evaluate global tRNAome alterations – four hours, 24 hours and five days, representing the immediate, intermediate and prolonged effect (**Fig. 2a**). Because the loss of TRMT6/61A noticeably impacted cell growth (**Fig. 1j**), cell confluency was carefully controlled to ensure all samples were collected at ∼ 70% confluency across conditions (see **Methods**). To overcome the interference of diverse tRNA modifications during reverse transcription, we optimized the NEB low bias small RNA-seq protocol by utilizing the NEB Induro reverse transcriptase^43^. This modification facilitated comprehensive tRNAome profiling, capturing mature tRNAs, precursor tRNAs and tRNA fragments (**Fig. 2a**). Our modified protocol successfully detected m^1^A_58_ at a misincorporation rate comparable to previously published tRNA-seq data for the same cell line^44^ (**Supplementary Fig. 2a**).

**Figure 2.**
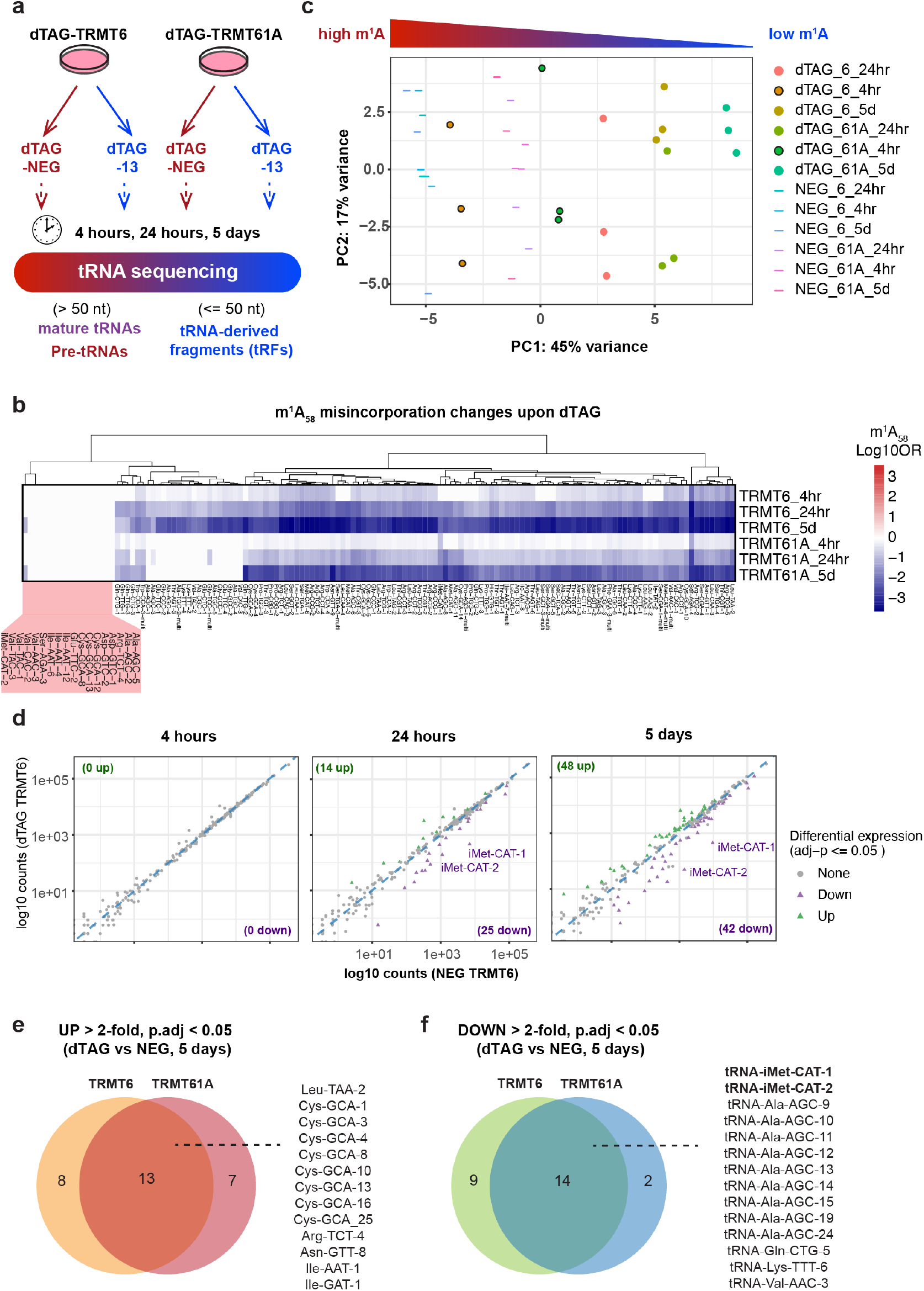
Loss of TRMT6/61A causes rapid global tRNAome remodeling. **(a)** Scheme of tRNA-sequencing strategy to profile global tRNA changes following the temporal loss of TRMT6/61A. **(b)** Heatmap showing m^1^A_58_ misincorporation change caused by dTAG at each time point, blue means decrease. Misincorporation rate at each time point is normalized to its respective dTAG-13-NEG control, and shown as Log 10 odds ratio. The red shades on the left highlight a group of tRNAs whose m^1^A_58_ misincorporation remains constant upon loss of TRMT6/61A. **(c)** PCA plot showing the cluster of all samples by mature tRNA expression at isodecoder levels. **(d)** Significantly differential tRNA isodecoders in HA-dTAG-TRMT6 HEK293T cells (adjusted p value <= 0.05). **(e-f)** Overlap of significantly up- or down-regulated tRNA isodecoders in HA-dTAG-TRMT6 and -TRMT61A HEK293T cells after 5 days of dTAG treatment (adjusted p value < 0.05, fold change > 2). See also **Supplementary Figure 2** and **Supplementary Table 3-8**.

Focusing on mature tRNAs, we observed a time-dependent decrease of m^1^A at position 58 across most tRNA families, which is consistent with the targeted degradation of the TRMT6/61A complex (**Fig. 2b** and **Supplementary Fig. 2b**). Surprisingly, while overall tRNA expression remained stable after the initial four hours of TRMT6/61A degradation, the reduction in m^1^A_58_ was already significant at this early time point (**Fig. 2b** and **Supplementary Fig. 2b**). Given the typically long half-lives of mature tRNAs over days^45^, this rapid decline suggests several possible mechanisms: a dilution effect driven by rapid transcription of new, unmodified tRNAs; the presence of active eraser activity; or the stress-induced tRNA cleavage of pre-existing, modified tRNAs. Interestingly, m^1^A levels on a small subset of tRNAs were insensitive to the dTAG treatment (**Fig. 2b**, red box; **Supplementary Table 1**). The stable m^1^A levels on these mature tRNAs suggest either they possess intrinsically low baseline m^1^A levels, or the hypomodified tRNAs are subject to active degradation, leaving only the modified pool detectable.

Following this initial 4-hour window, we evaluated the global tRNA expression changes at 24 hours and 5 days, and observed a robust sample clustering corresponding to m^1^A status (**Fig. 2c**), indicating significant remodeling of tRNA expression over time. Differential analysis across these later time points identified both down-regulated and up-regulated tRNAs at the isodecoder level (**Fig. 2d-f** and **Supplementary Fig. 2c-e**). Notably, tRNA^iMet-CAT-1/2^, tRNA^Ala-AGC-9-15/24^ and tRNA^Lys-TTT-6^ were consistently down-regulated after 24-hours and 5-days of dTAG treatment in both dTAG-TRMT6 and dTAG-TRMT61A cells, implying that the loss of m^1^A may destabilize these mature tRNAs. Conversely, tRNA-Leu-TAA-2 was consistently up-regulated after 24-hours of dTAG treatment, and a group of 13 tRNA isodecoders were up-regulated after 5-days of dTAG treatment. The up-regulated tRNAs upon TRMT6/61A degradation suggest a compensatory transcriptional response to maintain translational capacity; alternatively, it might be an unknown stabilization mechanism arising from competition between tRNA modification enzymes. A comprehensive list of differentially expressed tRNAs at each time point is provided in **Supplementary Table 3-8**. Altogether, the loss of TRMT6/61A induces rapid remodeling of the tRNAome at the mature tRNA level.

### Elongator but not initiator tRNAs are buffered by major isodecoders upon m^1^A loss

tRNAs are grouped by their anticodons into isoacceptor families, which are further composed of distinct tRNA genes known as isodecoders^45^. Despite a substantial number of tRNAs up- or down-regulated at the isodecoder level at five days (**Fig. 2**), less than ten tRNAs are up-regulated and down-regulated at the anticodon level at this time point (**Fig. 3a** and **Supplementary Fig. 3a**). The most significantly down-regulated tRNA was tRNA^iMet^, which comprises two isodecoder families. Both the major (tRNA-iMet-CAT-1) and minor (tRNA-iMet-CAT-2) isodecoders decreased progressively following dTAG treatment (**Fig. 3b**). In contrast to tRNA^iMet^, the loss of elongator tRNAs triggered by m^1^A loss appears to be buffered by their major isodecoders. For instance, while the minor isodecoder tRNA-Ser-AGA-4 decreased steadily by TRMT6 degradation (**Fig. 3c**), its major isodecoder counterpart, tRNA-Ser-AGA-2/1, remained stable (**Fig. 3d**). As a result, the overall abundance of tRNA-Ser-AGA at the anticodon level was unaltered. This highlights a buffering mechanism within the tRNAome and suggests that minor isodecoders may be intrinsically more sensitive to the m^1^A loss.

**Figure 3.**
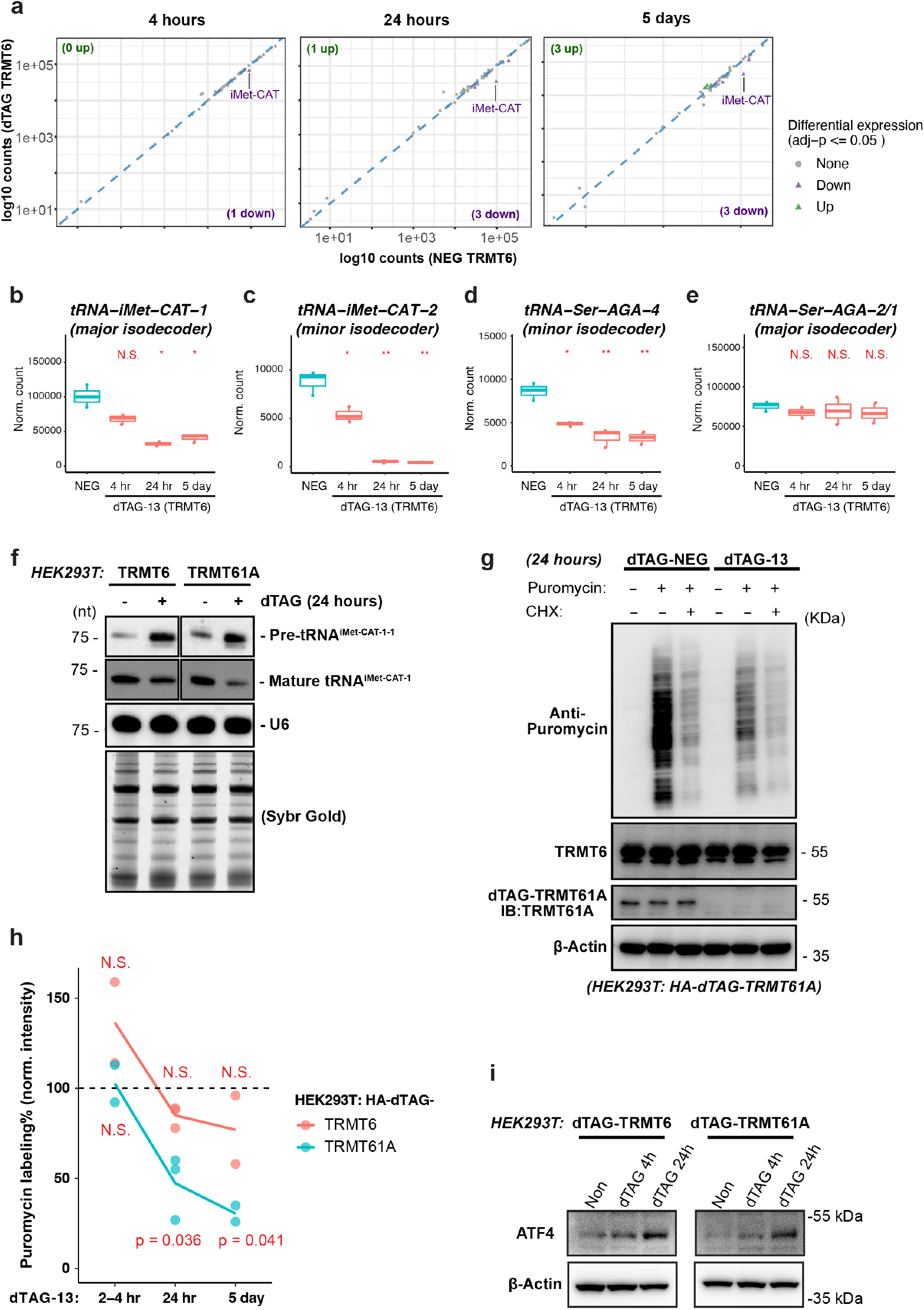
Elongator but not initiator tRNAs are buffered by major isodecoders upon m^1^A loss. **(a)** Significantly differential tRNAs summarized at anticodon level in HA-dTAG-TRMT6 HEK293T cells (adjusted p value <= 0.05). **(b-e)** Examples of tRNA isodecoder expression upon TRMT6 loss. tRNA-Ser-AGA but not tRNA-iMet-CAT loss is buffered by its major isodecoder expression. **(f)** Northern blot detection of mature and precursor tRNA^iMet-CAT^ upon 24-hours loss of TRMT6/61A in HEK293T. Mature tRNA^iMet-CAT^ is decreased and pre-tRNA^iMet-CAT^ is accumulated. **(g)** Example puromycin labeling to detect global protein synthesis change upon 24-hours loss of TRMT61A in HEK293T. Cycloheximide (CHX) treatment is included as a negative control. **(h)** Quantification of puromycin labeling upon temporal loss of TRMT6/61A in HEK293T (n = 2 or 3). **(i)** ATF4 western blot upon 24-hours loss of TRMT6/61A in HEK293T. Comparison between each condition and 100% were done by one-sample t test. See also **Supplementary Figure 3**.

The rapid depletion of mature initiator tRNA following TRMT6/61A loss was corroborated by Northern blot analysis in both HEK293T and HAP1 cells (**Fig. 3f** and **Supplementary Fig. 3b**), revealing a ∼50-80% reduction at the 24-hour time point. Because tRNA^iMet^ is strictly required for translation initiation and lacks a redundant major isodecoder to buffer its loss, we reasoned that its rapid depletion would act as a primary bottleneck for protein synthesis. Indeed, quantification via puromycin labeling demonstrated a robust decrease in global protein synthesis following 24 hours of TRMT6/61A loss and exacerbated over time (**Fig. 3g-h** and **Supplementary Fig. 3c-d**). This translational attenuation correlated with a rapid induction of ATF4, observable as early as four hours post TRMT6/61A degradation (**Fig. 3i** and **Supplementary Fig. 3c-d**). In summary, loss of TRMT6/61A selectively destabilizes initiator tRNAs, driving a rapid halt to global protein synthesis and activating ATF4 protein expression.

### Global accumulation of pre-tRNAs upon m^1^A loss suggests a defect in RNaseP/Z processing

Beyond the observed decrease in mature tRNA^iMet^, Northern blotting analysis of pre-tRNA^iMet-CAT-1-1^ revealed that the loss of TRMT6/61A leads to a dramatic accumulation of pre-tRNA^iMet^ species (**Fig. 3f** and **Supplementary Fig. 3b**). A similar effect on pre-tRNA^iMet^ accumulation has also been reported in *S. cerevisiae*^*46, 47*^. To determine whether this effect extends to other tRNAs, we re-analyzed our tRNA-seq data, specifically focusing on reads mapping exclusively to pre-tRNA sequences. A significant increase in pre-tRNA reads was observed as early as 4 hours post-dTAG treatment (**Fig. 4a-b** and **Supplementary Fig. 4a**). This accumulation is specific to the precursor species, as total read counts for mature tRNAs remained stable across all conditions (**Supplementary Fig. 4b-c**). In addition to pre-tRNA^iMet-CAT-1-1^ (**Fig. 3f**), we validated the accumulation of pre-tRNA^eMet-CAT-1-1^, pre-tRNA^Ile-TAT-2-3^ and pre-tRNA^Tyr-GTA-5-5^ following 24 hours of dTAG treatment (**Fig. 4c-d**). This processing defect is dependent on the complex’s catalytic activity: expression of wild-type TRMT61A, but not a catalytic inactive mutant (Q85A_D181A), rescued the accumulation caused by TRMT6/61A depletion (**Fig. 4e**). These data suggest that the observed increase in pre-tRNA levels stems from either upregulated transcription, or more likely, defective RNase P/Z processing during mature tRNA biogenesis.

**Figure 4.**
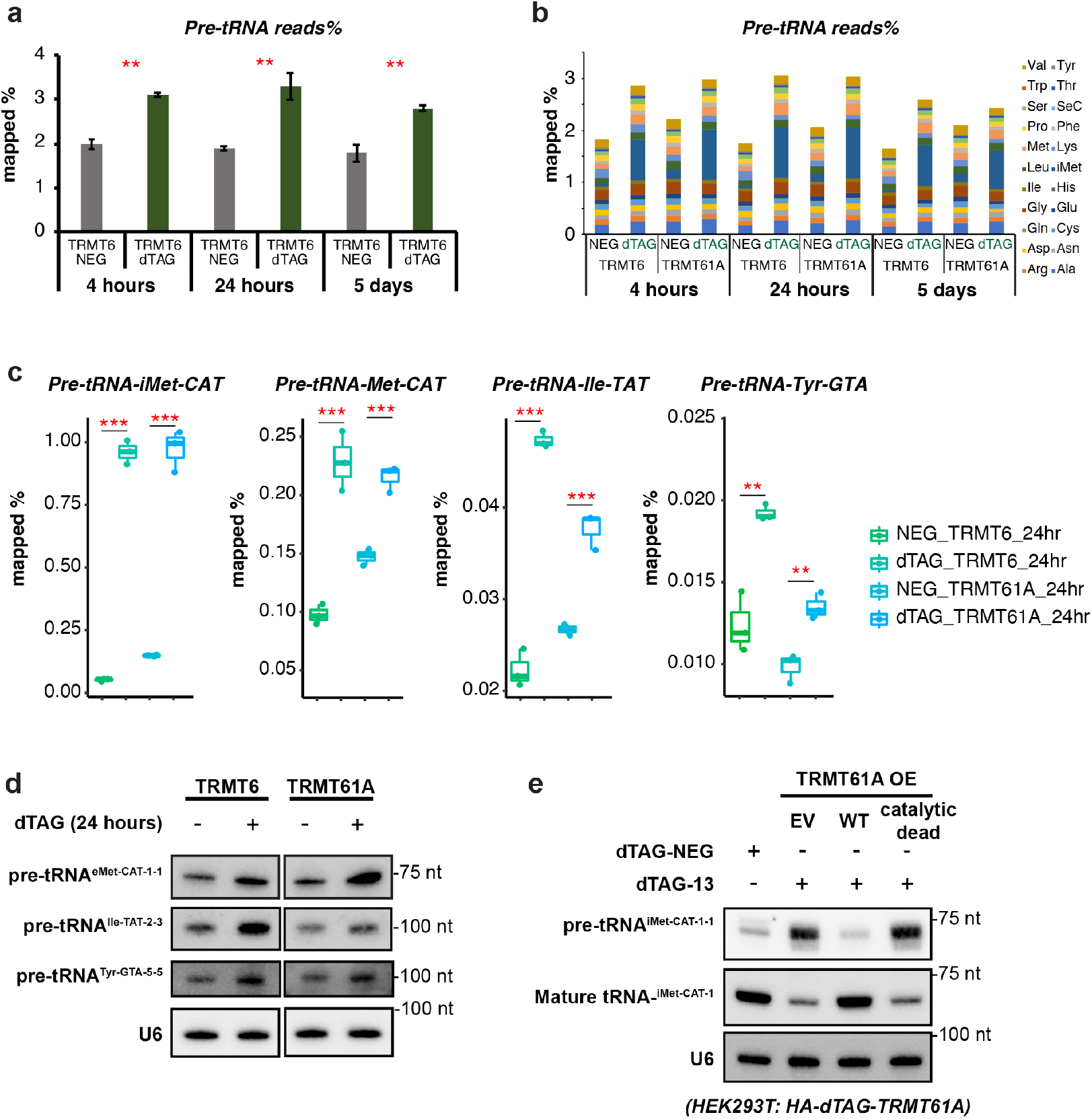
Global accumulation of pre-tRNAs upon m^1^A loss suggests a defect in RNaseP/Z processing. **(a-b)** Pre-tRNA-mapped reads are significantly increased upon TRMT6 degradation (n = 3). **(c)** Example pre-tRNAs that are significantly increased upon TRMT6/61A degradation. **(d)** Northern blot validation of individual pre-tRNAs (probe information see **Supplementary Table 1**). **(e)** Pre-tRNA-iMet-CAT-1-1 accumulation by TRMT61A loss can be rescued by over-expression (OE) of TRMT61A Wildtype (WT) but not the catalytic dead (Q85A_D181A). Comparison between two conditions were done by Student’s t test, two-tail; ** p < 0.01, *** p <0.001. See also **Supplementary Figure 4**.

### TRMT6/61A catalyzes m^1^A on pre-tRNAs and facilitates RNase Z processing

For our hypothesis to hold, m^1^A_58_ must be installed on pre-tRNAs prior to RNase P/Z processing. Historically, dedicated pre-tRNA sequencing datasets have been lacking because nascent tRNA transcripts harbor 5’ triphosphate (5’ ppp), which is poorly captured by standard RNA-seq protocols. To circumvent this, we pre-treated RNA samples with RppH to converts 5’ triphosphate to 5’ monophosphate prior to library preparation. As expected, RppH treatment markedly increased pre-tRNA coverage (**Supplementary Fig. 5a**). Crucially, we detected the characteristic m^1^A_58_ misincorporation signature on pre-tRNA reads containing trailer, leader, and even intron sequences (**Fig. 5a**).

**Figure 5.**
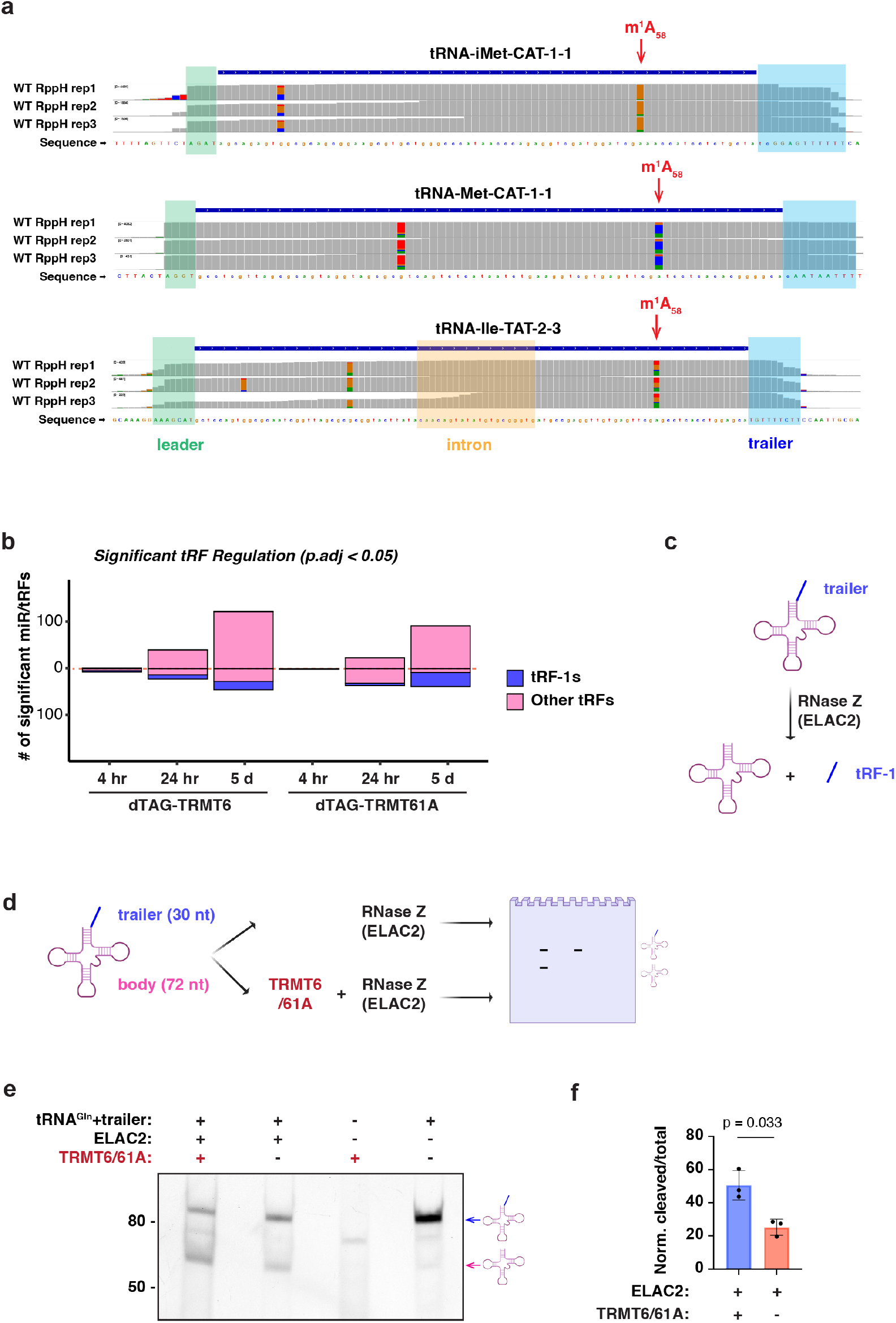
TRMT6/61A catalyzes m^1^A on pre-tRNAs and facilitates RNase Z processing. **(a)** IGV screenshots of m^1^A_58_ mismatch detected on Pre-tRNA-iMet-CAT-1-1, Pre-tRNA-Met-CAT-1-1 and Pre-tRNA-Ile-TAT-2-3. WT HEK293T total RNA was treated with RppH to increase capture of premature tRNA sequences. Green shade = leader sequences, Green shade = trailer sequences, Orange shade = intron sequences. **(b)** Number of significantly up- or down-regulated tRNA-derived fragments (p.adj < 0.05 by DESeq2). While tRF-1s are decreased by TRMT6/61A loss, other tRF types are up-regulated. **(c)** Schematic representation of tRF-1 generation by RNase Z (ELAC2) cleavage of the trailer sequence. **(d)** Scheme of *in vitro* RNase Z (ELAC2) assay with and without TRMT6/61A. **(e)** *In vitro* RNase Z assay product was resolved on TBE Urea gel to separate the substrate (top band) and the cleaved product (bottom band). **(f)** Quantification of in vitro RNase Z assay (n = 3, comparison done by paired two-tail t test). See also **Supplementary Figure 5**.

Building on this evidence of early m^1^A installation, we investigated the global landscape of tRNA-derived fragments (tRFs) across our temporal sequencing data. tRFs are categorized by their tRNA origins^48^. While most tRF types are derived from mature tRNAs (such as tRF-5s, tRF-3s, and tiRNAs), tRF-1s are specifically generated from the 3’ trailer sequences of pre-tRNAs as a direct byproduct of RNase Z cleavage^49^. We noticed an interesting dichotomy upon TRMT6/61A depletion (**Fig. 5b-c**): tRF-1s, specifically defined to start at the RNase Z cleavage site, were significantly decreased. In contrast, other tRF types derived from mature tRNAs were globally increased, likely reflecting canonical stress activation (**Supplementary Fig. 5b-d**). This transcriptome-wide depletion of tRF-1s strongly indicates a profound defect in RNase Z processing.

To directly assay whether TRMT6/61A facilitates processing, we performed *in vitro* RNase Z assay using tRNA substrate containing trailer sequence. The addition of recombinant TRMT6/61A significantly enhanced cleavage by the human RNase Z enzyme, ELAC2 (**Fig. 5d-e**). Collectively, these data indicate that TRMT6/61A-mediated m^1^A installation is an early tRNA modification event that is pivotal for downstream RNase Z processing, likely through promoting proper pre-tRNA folding.

### Hypomodified mature tRNA^iMet^ is degraded by XRN2, and XRN2 depletion rescues growth defects caused by TRMT6/61A loss

The rapid decrease in mature tRNA^iMet^ following TRMT6/61A depletion (**Fig. 3**) points to an active surveillance mechanism targeting hypomodified tRNA. In *S. cerevisiae*, both 5’-to-3’ and 3’-to-5’ decay pathways contribute to the elimination of hypomodified tRNA^iMet^ ^37, 38, 39, 40, 41^, but whether these parallel surveillance pathways are functionally conserved in mammalian cells remains poorly understood. To identify the responsible nucleases for this process, we knocked down the 5’-to-3’ exoribonucleases XRN1 and XRN2, the nuclear exosome complex catalytic subunits DIS3 (RRP44) and EXOSC10 (RRP6), and the poly(U)-binding protein SSB (Lupus La protein) in combination with TRMT6/61A degradation. Strikingly, only XRN2 knockdown fully rescued the dTAG-induced loss of tRNA^iMet^ (**Fig. 6a**), establishing XRN2 as the primary enzyme responsible for degrading hypomodified mature tRNA^iMet^ in human cells.

**Figure 6.**
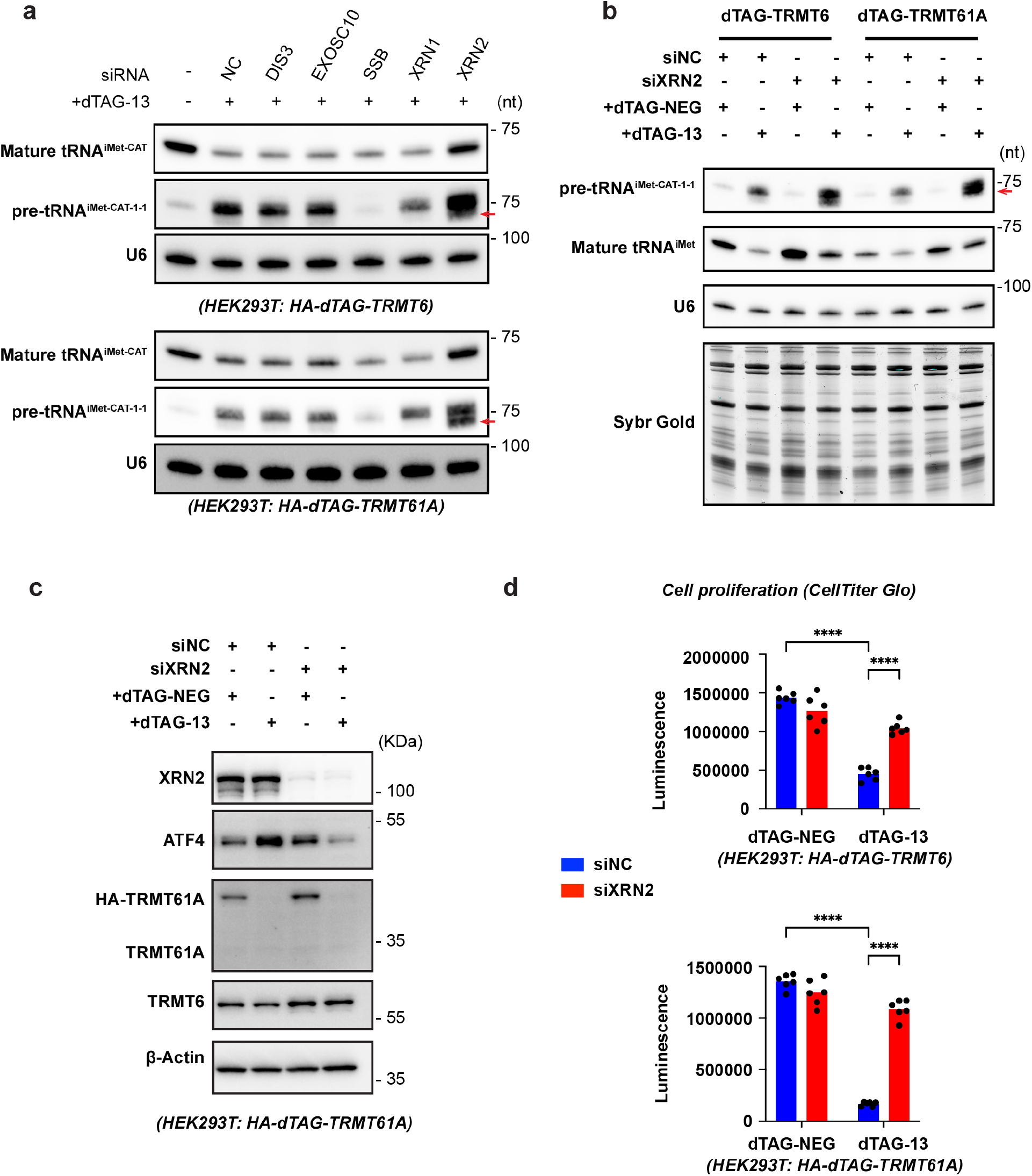
Hypomodified mature tRNA-iMet is degraded by XRN2, and XRN2 depletion rescues growth defects caused by TRMT6/61A loss. **(a)** Mature tRNA^iMet-CAT^ loss by TRMT6/61A degradation over five days is rescued by siXRN2 but not the other candidates, detected by Northern blot. **(b)** Mature and pre-tRNA^iMet-CAT^ changes by TRMT6/61A degradation is rescued by siXRN2. **(c)** ATF4 activation by TRMT61A degradation is rescued by siXRN2. **(d)** The decreased cell viability by TRMT6/61A degradation over five days is rescued by siXRN2, detected by CellTiter Glo assay (n = 6). See also **Supplementary Figure 6**.

Intriguingly, Northern blotting with a probe targeting the 3’ trailer sequence revealed the accumulation of a distinct product migrating just below pre-tRNA^iMet^ specifically upon combined siXRN2 and dTAG treatment (**Fig. 6a-b**, red arrow). This accumulation suggests that XRN2 also targets a hypomethylated, RNase P-cleaved precursor intermediate. This activity aligns with the biochemical requirement of XRN2, which demands a 5’ monophosphate group and is thus unable to act on the 5’ triphosphorylated initial transcript prior to RNase P processing.

In *S. cerevisiae*, hypomodified pre-tRNA^iMet^ is targeted by the TRAMP complex for poly-adenylation by TRF4 and subsequent 3’-to-5’ degradation by RRP44/DIS3^46^. While we detected baseline pre-tRNA^iMet^ transcripts in our tRNA-seq data, we did not observe an accumulation of polyadenylated pre-tRNA^iMet^ reads upon TRMT6/61A depletion alone, and DIS3 knockdown did not lead to accumulation of pre-tRNA^iMet^ (**Fig. 6b**). Although definitively ruling out the exosome pathway would require high-throughput RNA sequencing under concurrent DIS3 knockdown and dTAG depletion to capture potentially transient intermediates, our current data indicate that the 3’-to-5’ surveillance pathway is not the primary surveillance mechanism for hypomodified pre-tRNA^iMet^ in human cells.

Having established that XRN2 mediates the destruction of hypomodified tRNA^iMet^, we next asked whether blocking this degradation pathway could alleviate downstream cellular pathology. Remarkably, the rescue of mature tRNA^iMet^ levels by siXRN2 completely suppressed the downstream activation of the ATF4 stress response in dTAG-TRMT61A cells (**Fig. 6c**). Phenotypically, this molecular rescue translated into a dramatic restoration of cellular fitness. XRN2 knockdown significantly rescued the profound growth defects induced by loss of either TRMT6 or TRMT61A (**Fig. 6d**). To determine if this epistatic relationship represents a broader biological principle across human malignancies, we leveraged DepMap data encompassing over 1000 cancer cell lines. This pan-cancer analysis revealed a highly significant genetic co-dependency between XRN2 and both subunits of the methyltransferase complex, TRMT6 and TRMT61A (**Supplementary Fig. 6**). Taken together, these findings unveil a functional axis between XRN2 and TRMT6/61A machinery, demonstrating that XRN2-mediated clearance of hypomodified tRNA^iMet^ is the principal driver of the cellular pathology triggered by m^1^A_58_ deficiency.

## Discussion

In this study, we demonstrate that the TRMT6/61A complex and its associated m^1^A_58_ modification constitute as a critical checkpoint coordinating human tRNA maturation and translational homeostasis. Acute loss of TRMT6/61A using dTAG system swiftly remodels the human tRNAome (**Fig. 1-2**). While elongator tRNA fluctuations are buffered by isodecoder redundancy, the essential initiator tRNA^iMet^ is selectively depleted, driving a global decline in protein synthesis and activating the ATF4 translational stress response (**Fig. 3**). Our data reveal that m^1^A_58_ is installed early on precursor tRNAs, and facilitate efficient processing by RNase Z. In its absence, unprocessed pre-tRNAs aberrantly accumulate and tRNA-derived fragment (tRF) dynamics are heavily perturbed (**Fig. 4-5**). Finally, we identify the 5’-to-3’ exoribonuclease XRN2 as the executor of this quality control pathway. XRN2 actively degrades hypomodified mature tRNA^iMet^, and its inhibition rescues both tRNA^iMet^ abundance and stress-induced growth defects (**Fig. 6**).

Our findings reveal a novel and pivotal role of TRMT6/61A complex in facilitating pre-tRNA end processing. This phenomenon mirrors regulatory mechanisms observed in the mitochondria RNase P complex, where both the non-catalytic scaffolding role of the TRMT10C and its specific methyltransferase activity have been implicated in selectively facilitating RNase P cleavage for specific pre-tRNA substrates^50, 51^. Analogously, we hypothesize that the requirement for TRMT6/61A is highly likely to be tRNA-specific, supported by the observation that only a subset of pre-tRNAs accumulates upon TRMT6/61A loss. Because both RNase P and RNase Z (ELAC2) recognize the global 3D architecture of tRNA, especially the tRNA elbow, any structural disruption can impair processing. For instance, the human ELAC2 N-terminal domain features a ‘flexible arm’ that directly engages the T loop in pre-tRNA substate^52, 53^. The key location of m^1^A_58_ within the reverse Hoogsteen closing pair of the T-loop may allow it to facilitate and stabilize the D-T loop tertiary interactions by extending a methyl group ideally positioned to stack with the intercalating G18 nucleobase of the D-loop, introducing a positive charge to *N*1 atom that reinforces stacking with flanking nucleobases, and precluding A58 from unscheduled Watson-Crick-Franklin pairing with other nucleotides during tRNA folding. Additional structural and biophysical investigations will be needed to define and quantify the energetic contributions of m^1^A_58_ to tRNA folding and tertiary structure.

The physiological importance of this T-loop modification is underscored by recent work demonstrating that a loss of m^1^A_58_ on tRNA^Tyr^ in *S. cerevisiae* disrupts translation elongation, leading to ribosome collisions and triggering of the integrated stress response^54^. Intriguingly, TRMT6/61A deficiency appears to impact diverse tRNAs at the individual isodecoder level (**Fig. 2f**). While it has been suggested that m^1^A_58_ on mature tRNA^iMet^ is important for its proper folding to prevent exoribonucleolytic decay^55^, whether these downregulated elongator tRNAs (minor isodecoders) are similarly targeted via an XRN2-dependent mechanism remains to be determined.

Finally, our data indicate that human tRNA quality control diverges from *S. cerevisiae*, relying predominantly on XRN2 surveillance rather than the TRAMP complex to clear hypomodified tRNA^iMet^, a mechanism more closely resembling that of *S. pombe*^*41*^. The co-dependency between XRN2 and TRMT6/61A introduces profound clinical implications. For instance, malignancies overexpressing TRMT6/61A might present a unique therapeutic vulnerability for XRN2 inhibition. Conversely, developmental disorders characterized by mutant TRMT61A, such as Cornelia de Lange Syndrome^36^, might benefit from XRN2 targeting to stabilize compromised tRNA pools.

In summary, these findings support a model (**Fig. 7**) in which early m^1^A_58_ installation facilitates precursor tRNA end processing and protects mature tRNA^iMet^ from XRN2-mediated degradation to sustain cellular fitness. This framework links pathological variations in tRNA modifications to structural bottlenecks within the canonical tRNA maturation pathway.

**Figure 7.**
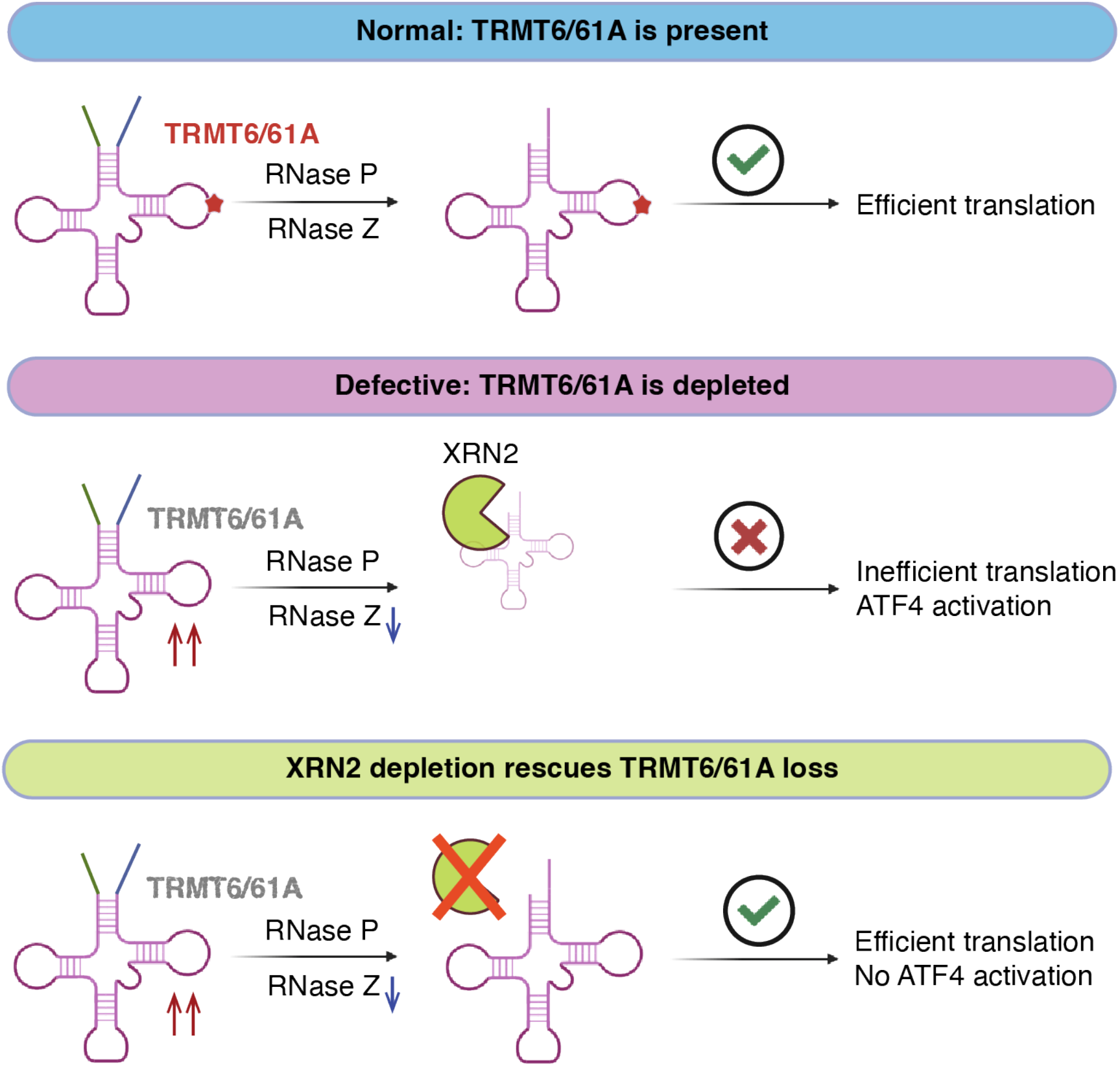
Proposed mechanism of TRMT6/61A-mediated tRNA processing and quality control by XRN2. Under normal conditions, the TRMT6/61A complex catalyzes the m^1^A modification (red star) on pre-tRNAs, facilitating proper processing by RNase P/Z. This ensures the production of stable mature tRNAs capable of supporting efficient translation. Upon TRMT6/61A depletion, the lack of early m1A installation impairs RNase Z processing, resulting in the accumulation of pre-tRNAs. The resulting hypomodified mature tRNA^iMet^ are recognized and rapidly degraded by the exoribonuclease XRN2, causing a global halt to translation and subsequent activation of ATF4. Co-depletion of XRN2 stabilizes hypomodified mature tRNA^iMet^, thereby rescuing translational capacity and prevent stress-induced ATF4 activation despite the upstream processing defects.

## Methods

### EXPERIMENTAL MODEL AND SUBJECT DETAILS

#### Cell culture

HEK293T (female) was maintained in HyClone Dulbecco’s High Glucose Modified Eagles medium with L-glutamine (Cyvita, #16750-074) plus 10% fetal bovine serum (Cyvita, #SH3091003) and 1% penicillin/streptomycin (Gibco, #15140122). HEK293T was obtained from ATCC (ATCC, #CRL-3216). HAP1 (male) was a kind gift from Wioletta Czaja (originally from Horizon Discovery), was maintained in IMDM (Iscove’s Modified Dulbecco’s Medium) with L-glutamine (Gibco, # 12440053) plus 10% fetal bovine serum and 1% penicillin/streptomycin. Mycoplasma contamination was routinely checked by mycostrip kit (InvivoGen, #rep-mysnc-50). Cells were grown in humidified incubators with 5% CO_2_ at 37°C.

#### Construction of HEK293T and HAP1 cell lines for dTAG-inducible protein degradation by CRISPR/Cas9

Degron tag (FKBP^F36V^)-containing knock-in cassettes were introduced into either the first or last coding exon of the TRMT6 or TRMT61A genes in HEK293T or HAP1 cells, yielding the N-terminal or C-terminal TRMT6-or TRMT61A-degradable cell lines, respectively, as previously described ^56^. Briefly, HEK293T or HAP1 cells were seeded in 10-cm plates and co-transfected with 6 µg of pCRIS-PITChv2-BSD-dTAG (TRMT6 or TRMT61A) donor plasmids (modified from Addgene #91792) and 6 µg of pX330A-nTRMT6 or -nTRMT61A/PITCh Cas9/gRNA plasmids (modified from Addgene #91794) using Lipofectamine 3000 (Invitrogen, #L3000015), according to the manufacturer’s instructions. 6 hours after transfection, appropriate fresh growth media was added to the cells for 48 hours. Cells were then selected with 10 µg/mL blasticidin, and single-cell clones were isolated in 96-well format. The donor and Cas9/gRNA plasmids are listed in **Supplementary Table 1**.

For genomic DNA isolation, cells were washed twice with cold PBS and lysed in 60 μL of QuickExtract DNA Extraction Solution (LGC, QE09050) by vortexing for 15 seconds. Samples were incubated at 65 °C for 6 minutes with shaking at 800 rpm, vortexed for 15 seconds, and then incubated at 98 °C for 2 minutes to inactivate proteinase K. Genotyping PCR was performed using 1 μL of genomic DNA, 2× Q5 hot start master mix (NEB), and primers listed in **Supplementary Table 1** to confirm correct knock-in as homozygous.

## METHOD DETAILS

### dTAG treatment and RNA collection

dTAG-13 (MilliporeSigma, #SML2601) and dTAG-13-NEG (TOCRIS, #6916) were dissolved in DMSO to make 1 mM stock. dTAG treatment was performed with indicated duration and concentrations, fresh media with compound was replenished after 3 days. For small RNA-seq experiment, cells were treated with dTAG-13 or dTAG-NEG at 500 nM final concentration. In order to keep the cell confluency about 70% before we harvest the cells, different numbers of cells were seeded depending on the treatment duration with either dTAG-13 or dTAG-NEG. For the dTAG-NEG treatment group, 6 × 10^6^ cells were seeded per 100-mm dish for the 4-hour treatment condition, 3 × 10^6^ cells per 100-mm dish for the 24-hour condition, and 5 × 10^4^ -1 × 10^5^ cells per 100-mm dish for the 5-day condition. And for the dTAG-13 treatment group, 6 × 10^6^ cells were seeded per 100-mm dish for the 4-hour treatment condition, 3 × 10^6^ cells per 100-mm dish for the 24-hour condition, and 5 × 10^5^ to 1 × 10^6^ cells per 100-mm dish for the 5-day condition. Total RNAs was extracted from cell lines by TRIzol Reagent (Invitrogen, #15596018) and Direct-zol RNA Miniprep Plus Kit with on-column DNase treatment (ZYMO, #R2071). We increased the EtOH:Trizol ratio to 2:1 (v/v) to retain small RNAs.

### Small RNA-seq library preparation

The small RNA-seq library was prepared with NEBNext Low-bias Small RNA Library Prep Kit (NEB, #E3420) with the following modifications. cDNA synthesis step was performed with Induro Reverse Transcriptase (NEB, #M0681) at 42 °C overnight to better overcome the RT-stalling modifications, and purified with 2X volume of NEBNext Sample Purification Beads to recover more tRNA-length inserts. After PCR with NEBNext LV Unique Dual Index Primers, the reaction was cleaned up with Purification Beads without size selection. Libraries were pooled and sequenced (PE150) on Illumina NovaSeq (Novogene). In general, each library has 10-20 million reads. To capture more pre-tRNA reads, wildtype HEK293T total RNA was pre-treated with RppH (NEB, #M0356) in Thermopol Reaction Buffer (NEB, #B9004S) at 37 °C for 1 hour, purified by SPRI beads before proceeding to small RNA-seq library preparation as above.

### Small RNA-seq data analysis

Cutadapt v4.9^57^ was used to trim 3’ adaptor sequence based on NEBNext kit (NEB, #E3420) and discard trimmed read length shorter than 15 nt. The adaptor-trimmed reads were first divided by the insert length - longer than 50 nt were analyzed for mature/pre-tRNAs and the remaining reads for tRF analysis. For mature tRNA quantification, adaptor-trimmed reads were analyzed by mim-tRNAseq v1.3.8 pipeline with the default setting (https://github.com/nedialkova-lab/mim-tRNAseq)^58^. The differentially expression results from DESeq2^59^ at isodecoder level are included in **Supplementary Table 3-8**. For precursor tRNA quantification, adaptor-trimmed reads were analyzed by tRAX (https://github.com/UCSC-LoweLab/tRAX)^60^ was used to map reads to precursor tRNAs.

tRNA fragments were quantified similarly as before^61^. Briefly, unitas v1.7.3^62^ with SeqMap v1.0.13 ^63^ was used to map the reads first to human sequence of miRBase Release 22^64^, then to other small RNA sequences including genomic tRNA database^65^, Ensembl Release 97 and SILVA rRNA database Release 132^66^. Unless otherwise specified (for example when testing 0 mismatch allowed in **Fig. 2c**), unitas setting (-species_miR_only –species homo_sapiens) was used to allow 2 non-templated 3’ nucleotides and 1 internal mismatch for miRNA mapping, and 1 mismatch and 0 insertion/deletion for tRNA fragments and other small RNA mapping (equivalent to –tail 2 –intmod 1 –mismatch 1 –insdel 0). microRNA reads were grouped by mature microRNA names, tRF raeds were grouped by parental tRNAs and tRF types (cut-off for tRNA halves is 30-nucleotide long); in the case of multi-mapping, a read was counted as fraction distributed equally to avoid duplicate counts. DESeq2 was used on the count matrix containing microRNAs and tRFs to identify differentially expressed RNAs for each comparison^59^.

### Northern blot and m^1^A immuno-Northern blot

Northern blot and immuno-Northern blot were performed similarly as before^67^. Briefly, 1-2 μg purified total RNA were resolved on 10% or 15% Novex TBE-Urea gel (Invitrogen, #EC6885BOX) and transferred to Amersham Hybond-N + nylon membrane (Cytiva, #RPN203B) by Trans-Blot SD semi-dry transfer apparatus (Bio-Rad). The membrane was cross-linked with 254 nm wavelength by UV Crosslinker (Fisher brand). **(1) Northern blot:** After UV crosslinking, the membrane was blocked by ExpressHyb Hybridization Solution (Takara Bio, #636833) and probed with 5’ biotinylated DNA probe (Integrated DNA Technologies, probe sequences see **Supplementary Table 1**) following the manufacturer’s instructions. Hybridized membrane was detected with Chemiluminescent Nucleic Acid Detection Module Kit (Thermo Fisher, #89880). **(2) m**^**1**^**A immuno Northern blot:** After crosslinking, the membrane was blocked with 3% non-fat milk in PBST and probed with 1:2000 m^1^A mouse antibody (MBL, #D3453) in 4 °C overnight. Goat-anti-mouse (Cell Signaling, #7076V, used at 1:5,000) HRP-linked secondary antibodies were used for detection.

### Protein extraction and western blot

Western blot assay was performed as described previously^67^. Whole-cell lysates were isolated from tissue samples or cultured cells using RIPA lysis buffer (Thermo Fisher, #8990) containing 1% protease inhibitor mixture (Millipore Sigma, #P8340), 10 mM Sodium Fluoride (NaF) (Millipore Sigma, #S6776), and 1 mM Sodium Orthovanadate (Na_3_VO_4_) (NEB, #P0758L). Equivalent amounts of protein were separated by 4-20% Tris-Glycine SDS-PAGE (Thermo Scientific, #XP04205), and protein expression was quantified by western blotting with primary antibodies (The primary antibodies were listed in **Supplementary Table 2**). Goat-anti-mouse (Cell Signaling, #7076V, used at 1:5,000) and goat-anti-rabbit (Cell Signaling, #7074V, used at 1:5,000) HRP-linked secondary antibodies were used for detection of the respective targets. Blots were developed using Immobilon Western Chemiluminescent HRP Substrate (Millipore Sigma, #WBKLS0500) on BioRad Chemi Doc MP Imager. Band signal intensities were obtained with BioRad ImageLab Software (v5.2.1) and the relative target protein levels normalized to β-actin.

### Puromycin labeling assay

For each group, three dishes of cells were prepared in parallel. Dish #1 serves as the vehicle control and should receive no treatment. Dish #2 to measure protein synthesis by puromycin labeling, cells were treated with 2 μg/mL puromycin in a CO_2_ incubator at 37 °C for 10 minutes. For Dish #3 (cycloheximide control), cells were pre-treated with 25 μM cycloheximide in a CO_2_ incubator at 37 °C for 5 minutes before the puromycin labeling. All cells were washed twice with cold DPBS and lysed with RIPA buffer supplemented with 1× protease inhibitor and phosphatase inhibitors (NaF, 10 μM; Na_2_VO_3_, 1 mM). Cell lysates were resolved by SDS-PAGE and analyzed by immunoblotting as described in the Western blotting section.

### HA immunofluorescence

To promote cell attachment, HEK293T cells were seeded onto 8-well chamber slides (Thermo Scientific, #154534) pre-coated with Poly-L-lysine solution (Millipore Sigma, #P4707-50ML). Briefly, 200 μL of 0.01% Poly-L-lysine solution was added to each well of an 8-well chamber slide, and the slide was gently rocked to ensure even coating of the culture surface. After incubation for 30 min at room temperature, the solution was aspirated, and the wells were rinsed twice with sterile tissue culture–grade water. The slides were then allowed to dry for at least 2 hours before cell seeding. A total of 5 × 10^4^ HEK293T cells were seeded into each well in 500 μL complete medium and cultured in a CO_2_ incubator for 2 days. Immunofluorescence staining was subsequently performed using the Immunofluorescence Application Solutions Kit (Cell Signaling Technology, #12727S) to detect TRMT6 or TRMT61A signals. Briefly, cells were fixed with 4% formaldehyde prepared in 1× Wash Buffer (200 μL per well). The cells were then blocked with Blocking Buffer (200 μL per well) for 60 min at room temperature. Cells were incubated overnight at 4°C with HA monoclonal primary antibody (Cell Signaling Technology, #3724S, 1:800 dilution). The following day, specimens were incubated for 1 hour at room temperature in the dark with fluorochrome-conjugated secondary antibody (Cell Signaling Technology #4412, 1:1000 dilution). Slides were mounted using antifade mounting medium containing DAPI (Vectashield Plus, #H-2000). For optimal results, slides were allowed to cure overnight at room temperature before imaging. Finally, fluorescence images were acquired using ZEISS LSM700 confocal microscope.

### TRMT6/61A co-immunoprecipitation (co-IP)

To confirm TRMT6 and TRMT61A protein-protein interaction in the engineered cell lines, co-IP was performed as follows. HEK293T cells were washed twice with cold 1× DPBS and lysed in ice-cold non-denaturing lysis buffer containing 10 mM Tris-HCl (pH 7.4), 150 mM NaCl, 1 mM EDTA, 1% Triton X-100, and 0.5% NP-40, supplemented with protease inhibitor cocktail and phosphatase inhibitors (1 mM Na_3_VO_4_ and 10 mM NaF). Cell lysates were incubated on ice for 30 min with intermittent mixing and clarified by centrifugation at 12,000 rpm for 10 minutes at 4 °C. Protein concentrations were determined using the Qubit Protein Broad Range Assay (Invitrogen, #A50668). For each immunoprecipitation reaction, equal amounts of protein lysate (∼2 mg) were mixed with 25 μL Pierce™ Anti-HA Magnetic Beads (Life Technologies, #88837) and incubated overnight at 4 °C with rotation. The beads were subsequently washed five times with ice-cold 1× Wash Buffer freshly prepared (50 mM Tris-HCl, pH 7.4, 150 mM NaCl, and 0.2% Triton X-100 detergent). Bound proteins were eluted by boiling at 95 °C for 5 minutes in 2× Sample Buffer containing 125 mM Tris-HCl (pH 6.8), 4% SDS, 20% (v/v) glycerol, and 0.004% bromophenol blue. Immunoprecipitated proteins and input lysates were resolved by SDS–PAGE and analyzed by immunoblotting as described in the Western blotting section.

### XRN2 knock-down by siRNA

To enhance knockdown efficiency, a two-round siRNA transfection strategy was employed in conjunction with dTAG treatment to simultaneously degrade TRMT6/61A. siRNA transfections were performed using Lipofectamine RNAiMAX Transfection Reagent (Invitrogen, #13778150) according to the manufacturer’s instructions, with a final siRNA concentration of 10 nM. siRNAs targeting the indicated genes (DIS3, EXOSC10, SSB, XRN1, and XRN2) or a non-targeting control (NC) ONTARGETplus smartpool siRNAs were obtained from Horizon Discovery/Dharmacon. The first transfection was performed using a reverse-transfection approach. Cells were incubated with the transfection mixture for 48 hours before a second round of transfection was performed using a forward-transfection method. Cells were subsequently incubated for an additional 48–72 h prior to downstream analyses. 500 nM dTAG-13 or dTAG-NEG was added at both rounds of transfection. For protein analysis, cells were washed with cold DPBS, lysed in RIPA buffer supplemented with protease and phosphatase inhibitors, and analyzed as described in the Western blotting section. For RNA analysis, cells were washed with cold DPBS and extract RNA by TRIzol reagent. Equal amounts of RNA were resolved by Novex UREA TBE Gels and analyzed by Northern blot by using the indicated probe. The sequences of all siRNAs used in this study are listed in **Supplementary Table 1**.

### TRMT61A over-expression

HEK293T dTAG cells were seeded in complete growth medium and cultured overnight to achieve ∼60–80% confluency at the time of transfection. Plasmid transfections were performed using Lipofectamine 3000 Transfection Reagent (Invitrogen, #L3000015) according to the manufacturer’s instructions. pcDNA-Flag-TRMT61A or empty pcDNA-Flag vector controls (1 μg /mL final concentration) were transfected. Cells were incubated with the transfection mixture for 48-72 h prior to downstream analyses. Overexpression efficiency was confirmed by immunoblotting. For protein analysis, cells were washed with cold DPBS and lysed in RIPA buffer supplemented with protease and phosphatase inhibitors. Equal amounts of protein lysates were resolved by SDS-PAGE and analyzed by immunoblotting using the indicated antibodies. For RNA analysis, cells were washed with cold DPBS and extract RNA by TRIzol reagent. Equal amounts of RNA were resolved by Novex UREA TBE Gels and analyzed by Northern blot by using the indicated probe. All plasmids used in this study are listed in **Supplementary Table 1**.

### Cell viability and confluency measurement

Cell viability was assessed using the CellTiter-Glo 3D Luminescent Cell Viability Assay (Promega, #G9683) according to the manufacturer’s instructions. Briefly, HEK293T dTAG cells were seeded into black opaque 96-well plates (Fisher Scientific, #CLS3603) at the indicated densities and allowed to adhere overnight in complete growth medium. For dTAG-TRMT6 (D2S5#17) cells, seed 3000 cells in each well for the NEG group and the dTAG-13 group. For dTAG-TRMT61A (D5S7#27) cells, seed 5000 cells in each well for the NEG group and the dTAG-13 group. Cells were subsequently treated with the indicated compounds or vehicle controls for the specified durations. At the endpoint of treatment, the CellTiter-Glo reagent was equilibrated to room temperature and added directly to each well at a 1:1 ratio with the culture medium. Plates were incubated on an orbital shaker for 10 minutes at the speed of 200 rpm to induce cell lysis and then further incubated at room temperature for 20 minunites to stabilize the luminescent signal. Luminescence, which correlates with intracellular ATP levels and viable cell number, was measured using Promega GloMax. All experiments were performed with at least six biological replicates and repeated independently at least twice.

Cell confluency was quantified using an automated macro in ImageJ software. Brightfield cell images were captured by MilliCell (Millipore Sigma, #MDCI10000) and converted to 8-bit grayscale. A variance filter (radius = 1 pixel) was applied to highlight regions of high cellular texture, and segmented into binary masks using the Triangle thresholding algorithm. To total percentage of the area covered by cells was then automatically measured from the binary masks. Three images were taken from each time point.

### *In vitro* methylation and RNase Z assay

The expression plasmid, pET17b-6xHis-Trm6-Trm61, was kindly provided by Finer-Moore^68^, containing the 3C protease cleavage sequence C-terminal to the 6×His tag. The TRMT6/61A complex was expressed and purified as previously described^68, 69^. Human ELAC2 and tRNA^Gln^ + trailer substrate were prepared as previously^70^. For *in vitro* methylation and RNase Z assay, 20 μM tRNA was first folded in a buffer containing 25 mM Tris-HCl (pH 7.5) and 150 mM NaCl. The folding of 20 μM tRNA in a buffer containing 25 mM Tris-HCl (pH 7.5) and 150 mM NaCl was performed by first denaturing the tRNA at 90°C for 2 minutes, then rapidly cooled on ice. 10 mM MgCl2 was added to the tRNA to facilitate refolding. 1 μM folded tRNA was incubated with 0.1875 μM ELAC2 and 0.2 μM TRMT6/61A in buffer containing 25 mM Tris-HCl, pH. 7.5, 100 mM NaCl, 10 mM MgCl_2_, 1 mM SAM and 1 mM DTT at 37 °C for 1 hour. The reactions were stopped by adding 10 μL of Novex TBE-Urea Sample Buffer (2X) (Invitrogen, #LC6876), and the reactions were run on a 10% Tris-Borate-EDTA (TBE) Urea gel (Invitrogen, #EC68752). Gel signal intensities were quantified by Bio-Rad Image Lab, and three independent replicates were performed.

## QUANTIFICATION AND STATISTICAL ANALYSIS

Number of independent biological replicates and statistical details are described in the corresponding figure legends. Student’s t-test and Wilcoxon test were performed by Prism v10.4.1 or R v4.2.1 (“t.test” and “wilcox.test”). Differential analysis for small RNA-seq and RNA-seq was performed by DESeq2 (v1.36) Wald test with Benjamini-Hochberg adjustment for multiple hypothesis testing.

## DATA AVAILABILITY

The small RNA-seq generated in this study has been deposited in the Gene Expression Omnibus (GEO) database under accession code XXXXXX.

## ACKNOWLEDGEMENT

This research was supported by UAB Pittman Scholar fund (to Z.S.), NIH grants R00 CA259526 (to Z.S.), R35 ES031707 (to Y.W.), T32 GM146611 (to N.N.), T32 NS121721 (to M.A.), T32 GM135028 (to K.M.) and T32 ES018827 (to A.H.K.). We would like to thank Janet Finer-Moore (University of California at San Francisco) for sharing the TRMT6/61A expression plasmid. We would like to thank Coral Wille’s lab (University of Alabama at Birmingham) for sharing the dTAG plasmids and cloning protocol, Wioletta Czaja’s lab (University of Alabama at Birmingham) for sharing HAP1 cells, and Anindya Dutta’s lab (University of Alabama at Birmingham) for sharing antibodies. We would like to thank IT Research Computing Team at the University of Alabama at Birmingham for NGS support. Figure 7 was made by BioRender.

## AUTHOR CONTRIBUTIONS

Conceptualization, X.L. and Z.S.; Methodology, Investigation, Data Curation and Visualization, X.L., Z.S., N.N., K.M. and M.A.; Validation, A.J. and N.R.; Resources, X.L., Z.S., A.H.K., Y.W., I.S. and J.Z.; Writing – Original Draft, Z.S. and X.L.; Writing – Review & Editing, X.L., N.N., A.J., I.S., M.A., K.M., N.R., A.H.K., Y.W., J.Z., Z.S.; Funding Acquisition, Z.S., N.N., M.A., K.M. and Y.W.; Supervision, Z.S.

## REFERENCES

1. Phizicky EM, Hopper AK. The life and times of a tRNA. RNA 29, 898–957 (2023).

2. Orellana EA, Siegal E, Gregory RI. tRNA dysregulation and disease. Nat Rev Genet 23, 651–664 (2022).

3. Kirchner S, Ignatova Z. Emerging roles of tRNA in adaptive translation, signalling dynamics and disease. Nat Rev Genet 16, 98–112 (2015).

4. Zhang W, Foo M, Eren AM, Pan T. tRNA modification dynamics from individual organisms to metaepitranscriptomics of microbiomes. Mol Cell 82, 891–906 (2022).

5. Schaefer M, Kapoor U, Jantsch MF. Understanding RNA modifications: the promises and technological bottlenecks of the ‘epitranscriptome’. Open Biol 7, (2017).

6. Cayir A. RNA modifications as emerging therapeutic targets. Wiley Interdiscip Rev RNA 13, e1702 (2022).

7. Delaunay S, Helm M, Frye M. RNA modifications in physiology and disease: towards clinical applications. Nat Rev Genet 25, 104–122 (2024).

8. Cui L, et al. RNA modifications: importance in immune cell biology and related diseases. Signal Transduct Target Ther 7, 334 (2022).

9. Gilbert WV, Nachtergaele S. mRNA Regulation by RNA Modifications. Annu Rev Biochem 92, 175–198 (2023).

10. Boo SH, Kim YK. The emerging role of RNA modifications in the regulation of mRNA stability. Exp Mol Med 52, 400–408 (2020).

11. Sun H, Li K, Liu C, Yi C. Regulation and functions of non-m(6)A mRNA modifications. Nat Rev Mol Cell Biol 24, 714–731 (2023).

12. Barbieri I, Kouzarides T. Role of RNA modifications in cancer. Nat Rev Cancer 20, 303–322 (2020).

13. Begik O, Lucas MC, Liu H, Ramirez JM, Mattick JS, Novoa EM. Integrative analyses of the RNA modification machinery reveal tissue-and cancer-specific signatures. Genome Biol 21, 97 (2020).

14. Chujo T, Tomizawa K. Human transfer RNA modopathies: diseases caused by aberrations in transfer RNA modifications. FEBS J 288, 7096–7122 (2021).

15. Zhang K, Lentini JM, Prevost CT, Hashem MO, Alkuraya FS, Fu D. An intellectual disability-associated missense variant in TRMT1 impairs tRNA modification and reconstitution of enzymatic activity. Hum Mutat 41, 600–607 (2020).

16. Blaesius K, et al. Mutations in the tRNA methyltransferase 1 gene TRMT1 cause congenital microcephaly, isolated inferior vermian hypoplasia and cystic leukomalacia in addition to intellectual disability. Am J Med Genet A 176, 2517–2521 (2018).

17. Katsara O, Schneider RJ. m(7)G tRNA modification reveals new secrets in the translational regulation of cancer development. Mol Cell 81, 3243–3245 (2021).

18. Orellana EA, et al. METTL1-mediated m(7)G modification of Arg-TCT tRNA drives oncogenic transformation. Mol Cell 81, 3323–3338 e3314 (2021).

19. Nguyen NYT, Liu X, Dutta A, Su Z. The Secret Life of N(1)-methyladenosine: A Review on its Regulatory Functions. J Mol Biol 437, 169099 (2025).

20. Zhou H, et al. m1A and m1G disrupt A-RNA structure through the intrinsic instability of Hoogsteen base pairs. Nature Structural & Molecular Biology 23, 803–810 (2016).

21. Roundtree IA, Evans ME, Pan T, He C. Dynamic RNA Modifications in Gene Expression Regulation. Cell 169, 1187–1200 (2017).

22. Schwartz MH, et al. Microbiome characterization by high-throughput transfer RNA sequencing and modification analysis. Nat Commun 9, 5353 (2018).

23. Li X, et al. Base-Resolution Mapping Reveals Distinct m(1)A Methylome in Nuclear-and Mitochondrial-Encoded Transcripts. Mol Cell 68, 993–1005 e1009 (2017).

24. Dominissini D, et al. The dynamic N(1)-methyladenosine methylome in eukaryotic messenger RNA. Nature 530, 441–446 (2016).

25. Liu F, et al. ALKBH1-Mediated tRNA Demethylation Regulates Translation. Cell 167, 1897 (2016).

26. Richter U, et al. RNA modification landscape of the human mitochondrial tRNA(Lys) regulates protein synthesis. Nat Commun 9, 3966 (2018).

27. Zhang C, et al. The landscape of m(1)A modification and its posttranscriptional regulatory functions in primary neurons. Elife 12, (2023).

28. Chen Z, et al. Transfer RNA demethylase ALKBH3 promotes cancer progression via induction of tRNA-derived small RNAs. Nucleic Acids Res 47, 2533–2545 (2019).

29. Wei J, et al. Differential m(6)A, m(6)A(m), and m(1)A Demethylation Mediated by FTO in the Cell Nucleus and Cytoplasm. Mol Cell 71, 973–985 e975 (2018).

30. Wang Y, et al. N(1)-methyladenosine methylation in tRNA drives liver tumourigenesis by regulating cholesterol metabolism. Nat Commun 12, 6314 (2021).

31. Saikia M, Fu Y, Pavon-Eternod M, He C, Pan T. Genome-wide analysis of N1-methyl-adenosine modification in human tRNAs. RNA 16, 1317–1327 (2010).

32. Macari F, et al. TRM6/61 connects PKCalpha with translational control through tRNAi(Met) stabilization: impact on tumorigenesis. Oncogene 35, 1785–1796 (2016).

33. Zuo H, Wu A, Wang M, Hong L, Wang H. tRNA m(1)A modification regulate HSC maintenance and self-renewal via mTORC1 signaling. Nat Commun 15, 5706 (2024).

34. Liu Y, et al. tRNA-m(1)A modification promotes T cell expansion via efficient MYC protein synthesis. Nat Immunol 23, 1433–1444 (2022).

35. Tao EW, et al. TRMT6-mediated tRNA m(1)A modification acts as a translational checkpoint of histone synthesis and facilitates colorectal cancer progression. Nat Cancer 6, 1458–1476 (2025).

36. Landrum MJ, et al. ClinVar: public archive of relationships among sequence variation and human phenotype. Nucleic Acids Res 42, D980–985 (2014).

37. Zhang C, Jia G. Reversible RNA ModificationN1-Methyladenosine (m1A) in mRNA and tRNA. Genomics, Proteomics & Bioinformatics 16, 155–161 (2018).

38. Kawai G, et al. Conformational rigidity of specific pyrimidine residues in tRNA arises from posttranscriptional modifications that enhance steric interaction between the base and the 2’-hydroxyl group. Biochemistry 31, 1040–1046 (1992).

39. Grosjean H, Edqvist J, Stråby KB, Giegé R. Enzymatic Formation of Modified Nucleosides in tRNA: Dependence on tRNA Architecture. Journal of Molecular Biology 255, 67–85 (1996).

40. Anderson J, et al. The essential Gcd10p–Gcd14p nuclear complex is required for 1-methyladenosine modification and maturation of initiator methionyl-tRNA. Genes & Development 12, 3650–3662 (1998).

41. Tasak M, Phizicky EM. Initiator tRNA lacking 1-methyladenosine is targeted by the rapid tRNA decay pathway in evolutionarily distant yeast species. PLoS Genet 18, e1010215 (2022).

42. Nabet B, et al. The dTAG system for immediate and target-specific protein degradation. Nat Chem Biol 14, 431–441 (2018).

43. Nakano Y, et al. Genome-wide profiling of tRNA modifications by Induro-tRNAseq reveals coordinated changes. Nat Commun 16, 1047 (2025).

44. Behrens A, Rodschinka G, Nedialkova DD. High-resolution quantitative profiling of tRNA abundance and modification status in eukaryotes by mim-tRNAseq. Mol Cell 81, 1802–1815 e1807 (2021).

45. Pan T. Modifications and functional genomics of human transfer RNA. Cell Res 28, 395–404 (2018).

46. Kadaba S, Krueger A, Trice T, Krecic AM, Hinnebusch AG, Anderson J. Nuclear surveillance and degradation of hypomodified initiator tRNAMet in S. cerevisiae. Genes Dev 18, 1227–1240 (2004).

47. Anderson J, et al. The essential Gcd10p-Gcd14p nuclear complex is required for 1-methyladenosine modification and maturation of initiator methionyl-tRNA. Genes Dev 12, 3650–3662 (1998).

48. Su Z, Wilson B, Kumar P, Dutta A. Noncanonical Roles of tRNAs: tRNA Fragments and Beyond. Annu Rev Genet 54, 47–69 (2020).

49. Lee YS, Shibata Y, Malhotra A, Dutta A. A novel class of small RNAs: tRNA-derived RNA fragments (tRFs). Genes Dev 23, 2639–2649 (2009).

50. Karasik A, Fierke CA, Koutmos M. Interplay between substrate recognition, 5’ end tRNA processing and methylation activity of human mitochondrial RNase P. RNA 25, 1646–1660 (2019).

51. Vilardo E, Nachbagauer C, Buzet A, Taschner A, Holzmann J, Rossmanith W. A subcomplex of human mitochondrial RNase P is a bifunctional methyltransferase--extensive moonlighting in mitochondrial tRNA biogenesis. Nucleic Acids Res 40, 11583–11593 (2012).

52. Bhatta A, et al. Molecular basis of human nuclear and mitochondrial tRNA 3’ processing. Nat Struct Mol Biol 32, 613–624 (2025).

53. Li de la Sierra-Gallay I, Pellegrini O, Condon C. Structural basis for substrate binding, cleavage and allostery in the tRNA maturase RNase Z. Nature 433, 657–661 (2005).

54. Stuart CJ, Hurtig JE, Tzadikario T, Thomas NK, Jain M, van Hoof A. The highly conserved intron of tyrosine tRNA is critical for (m1)A58 modification and controls the integrated stress response. Proc Natl Acad Sci U S A 122, e2502364122 (2025).

55. Yared MJ, Yoluc Y, Catala M, Tisne C, Kaiser S, Barraud P. Different modification pathways for m1A58 incorporation in yeast elongator and initiator tRNAs. Nucleic Acids Res 51, 10653–10667 (2023).

56. Nabet B, et al. The dTAG system for immediate and target-specific protein degradation. Nature Chemical Biology 14, 431–441 (2018).

57. Martin M. Cutadapt removes adapter sequences from high-throughput sequencing reads. EMBnet J 17, 10–12 (2011).

58. Behrens A, Nedialkova DD. Experimental and computational workflow for the analysis of tRNA pools from eukaryotic cells by mim-tRNAseq. STAR Protoc 3, 101579 (2022).

59. Love MI, Huber W, Anders S. Moderated estimation of fold change and dispersion for RNA-seq data with DESeq2. Genome Biol 15, 550 (2014).

60. Chan PP, Holmes AD, Lowe TM. Analyzing, visualizing, and annotating tRNA-derived RNAs using tRAX and tDRnamer. Methods Enzymol 711, 103–133 (2025).

61. Su Z, et al. TRMT6/61A-dependent base methylation of tRNA-derived fragments regulates gene-silencing activity and the unfolded protein response in bladder cancer. Nat Commun 13, 2165 (2022).

62. Gebert D, Hewel C, Rosenkranz D. unitas: the universal tool for annotation of small RNAs. BMC Genomics 18, 644 (2017).

63. Jiang H, Wong WH. SeqMap: mapping massive amount of oligonucleotides to the genome. Bioinformatics 24, 2395–2396 (2008).

64. Kozomara A, Birgaoanu M, Griffiths-Jones S. miRBase: from microRNA sequences to function. Nucleic Acids Res 47, D155–D162 (2019).

65. Chan PP, Lowe TM. GtRNAdb 2.0: an expanded database of transfer RNA genes identified in complete and draft genomes. Nucleic Acids Res 44, D184–189 (2016).

66. Quast C, et al. The SILVA ribosomal RNA gene database project: improved data processing and web-based tools. Nucleic Acids Res 41, D590–596 (2013).

67. Su Z, et al. TRMT6/61A-dependent base methylation of tRNA-derived fragments regulates gene-silencing activity and the unfolded protein response in bladder cancer. Nature Communications 13, 2165 (2022).

68. Finer-Moore J, Czudnochowski N, O’Connell JD, 3rd, Wang AL, Stroud RM. Crystal Structure of the Human tRNA m(1)A58 Methyltransferase-tRNA(3)(Lys) Complex: Refolding of Substrate tRNA Allows Access to the Methylation Target. J Mol Biol 427, 3862–3876 (2015).

69. Sun Y, et al. m(1)A in CAG repeat RNA binds to TDP-43 and induces neurodegeneration. Nature 623, 580–587 (2023).

70. Skeparnias I, et al. Structural basis of MALAT1 RNA maturation and mascRNA biogenesis. Nat Struct Mol Biol 31, 1655–1668 (2024).

