## Supplementary Fig for "TRMT6/61A-mediated m^1^A methylation facilitates human pre-tRNA maturation and prevents surveillance by XRN2"

**Supplementary Figure legends**

**Supplementary Figure 1. Rapid degradation of endogenous TRMT6/61A by dTAG (related to Figure 1).**

**(a)** Co-IP to confirm dTAG does not interfere with TRMT6/61A complex formation. **(b)** Time course of 0.5 μM dTAG treatment in HA-dTAG-TRMT61A HEK293T cells. **(c)** Genomic PCR to confirm successful knock in in HAP1 cells. **(d-e)** Time course of 0.5 μM dTAG treatment in HA-dTAG-TRMT6/61A HAP1 cells. **(f-g)** Representative images of HEK293T HA-dTA-TRMT6/61A cells over the course of 5-days treatment with 0.5 μM dTAG-13 or dTAG-NEG treatment.

**Supplementary Figure 2. Loss of TRMT6/61A causes rapid global tRNAome remodeling (related to Figure 2).**

**(a)** Correlation of $m^1A_{58}$ misincorporation on cytosolic tRNAs in our HEK293T dataset and published mim-tRNA-seq dataset. **(b)** $m^1A_{58}$ misincorporation change on tRNA-iMet-CAT-1 caused by dTAG at each time point (n = 3). Misincorporation rate at each time point is compared to its respective dTAG-13-NEG control, and statistics was done by student's t test. **(c)** Significantly differential tRNA isodecoders in HA-dTAG-TRMT61A HEK293T cells (adjusted p value <= 0.05). **(d-e)** Overlap of significantly up- or down-regulated tRNA isodecoders in HA-dTAG-TRMT6 and -TRMT61A HEK293T cells after 24 hours of dTAG treatment (adjusted p value < 0.05, fold change > 2).

**Supplementary Figure 3. Elongator but not initiator tRNAs are buffered by major isodecoders upon $m^1A$ loss (related to Figure 3).**

**(a)** Significantly differential tRNAs summarized at anticodon level in HA-dTAG-TRMT61A HEK293T cells (adjusted p value <= 0.05). **(b)** Northern blot detection of mature and precursor $tRNA^{iMet\text{-}CAT}$ upon 24-hours loss of TRMT6/61A in HAP1. Mature $tRNA^{iMet\text{-}CAT}$ is decreased and pre- $tRNA^{iMet\text{-}CAT}$ is accumulated. **(c-d)** Puromycin labeling to detect global protein synthesis change and ATF4 western blot upon loss of TRMT6/61A in HAP1. Cycloheximide (CHX) treatment is included as a control for puromycin labeling.

**Supplementary Figure 4. Global accumulation of pre-tRNAs upon $m^1A$ loss suggests a defect in RNaseP/Z processing (related to Figure 4).**

**(a)** Pre-tRNA-mapped reads are significantly increased upon TRMT61A degradation (n = 3). **(b-c)** Mature tRNA-mapped reads are unchanged upon TRMT6/61A degradation (n = 3).
Comparison between two conditions were done by Student's t test, two-tail; ** p < 0.01, *** p <0.001.

**Supplementary Figure 5. TRMT6/61A catalyzes $m^1A$ on pre-tRNAs and facilitates RNase Z processing (related to Figure 5).**

**(a)** Read distribution suggests pre-tRNA coverage is increased with RppH treatment. **(b)** tRF reads are significantly increased upon 5 days of TRMT6/61A degradation (n = 3). **(c-d)** Box plot showing global small RNA changes divided by RNA types in dTAG-TRMT61A HEK293T cells after dTAG treatment for 5 days. Comparison between microRNA and each tRF group (divided by tRF type or amino acid group) is done by Wilcoxon test.

**Supplementary Figure 6. Hypomodified mature $tRNA^{iMet}$ is degraded by XRN2, and XRN2 depletion rescues growth defects caused by TRMT6/61A loss (related to Figure 6).**

**(a-b)** Co-dependency of TRMT6/61A with XRN2 is significant in DepMap CRISPR assay (Pearson correlation between Chronos Scores and p value is shown, n = 1186 cell lines). **(c)** Co-dependency between TRMT6 and TRMT61A is shown as positive control.

**Supplementary Figure 1.**

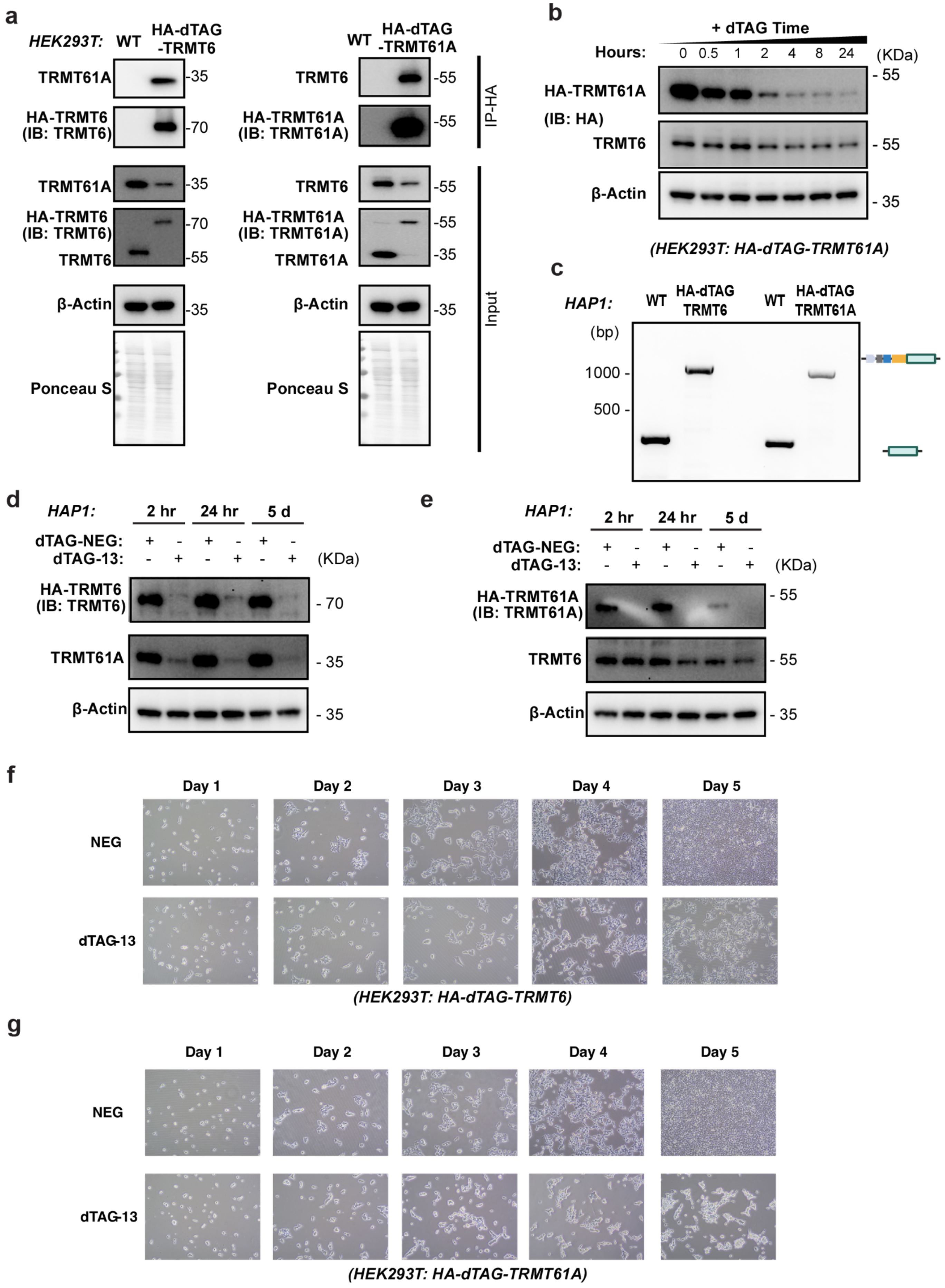

**Supplementary Figure 2.**

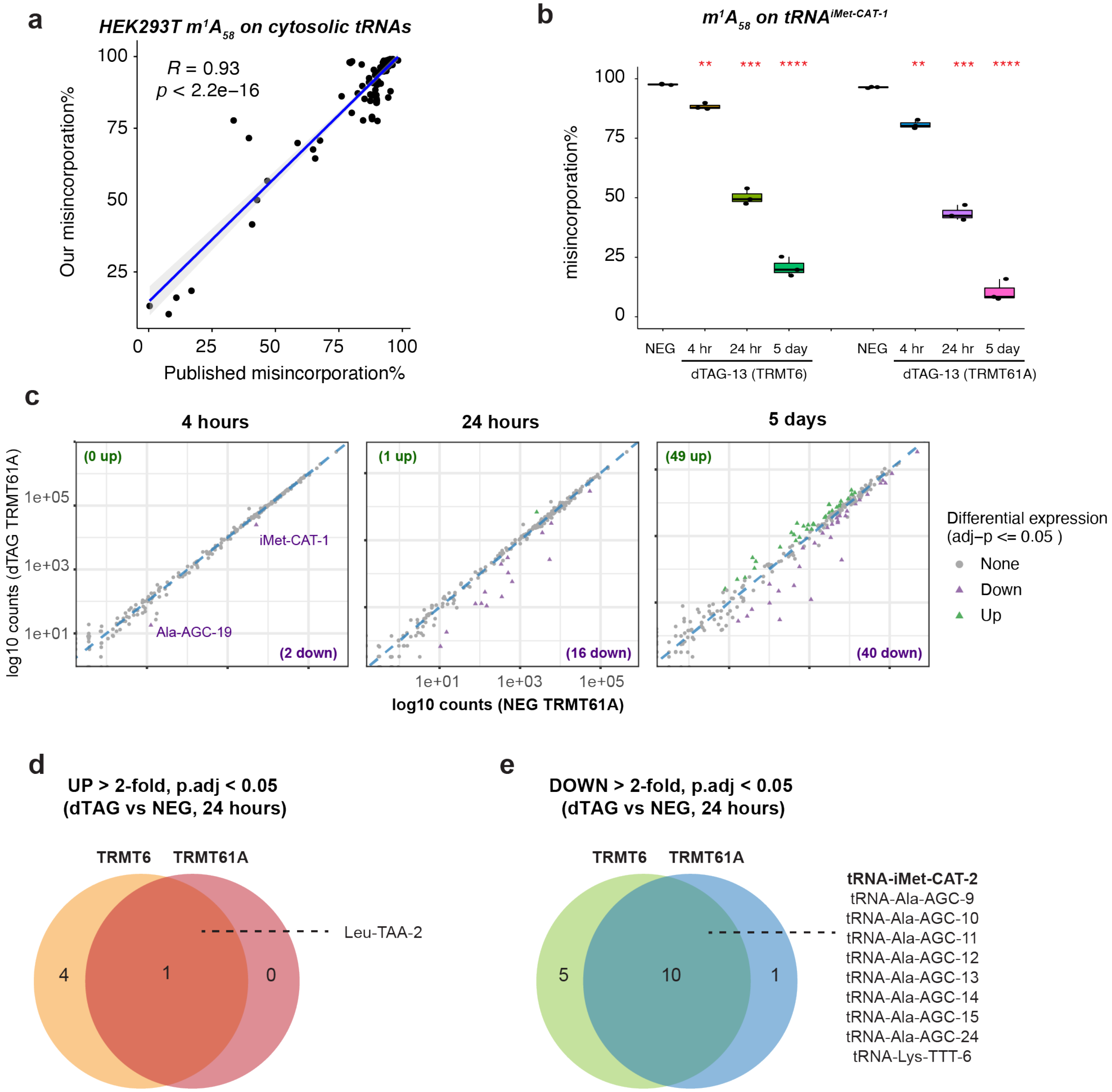

**Supplementary Figure 3.**

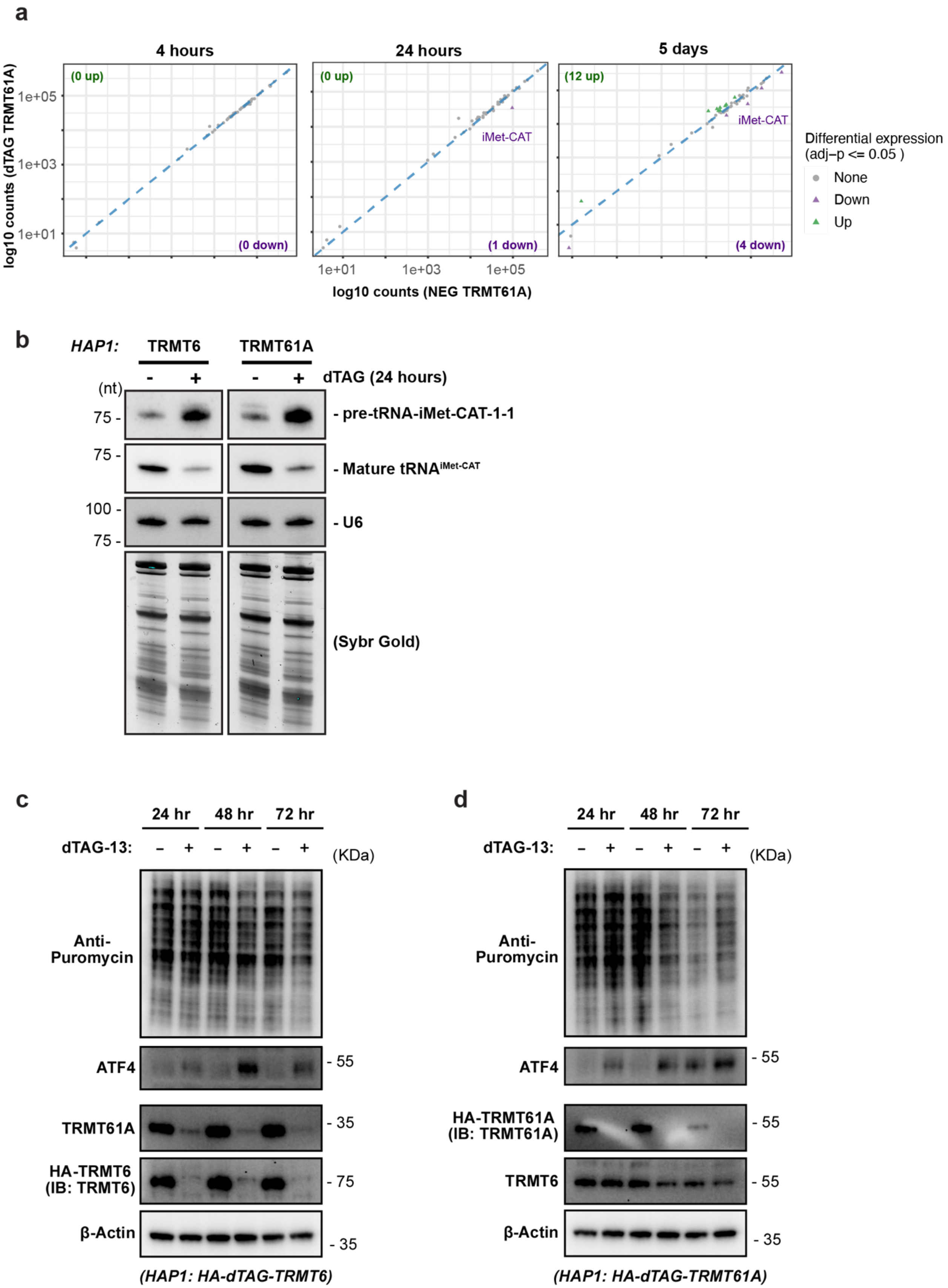

**Supplementary Figure 4.**

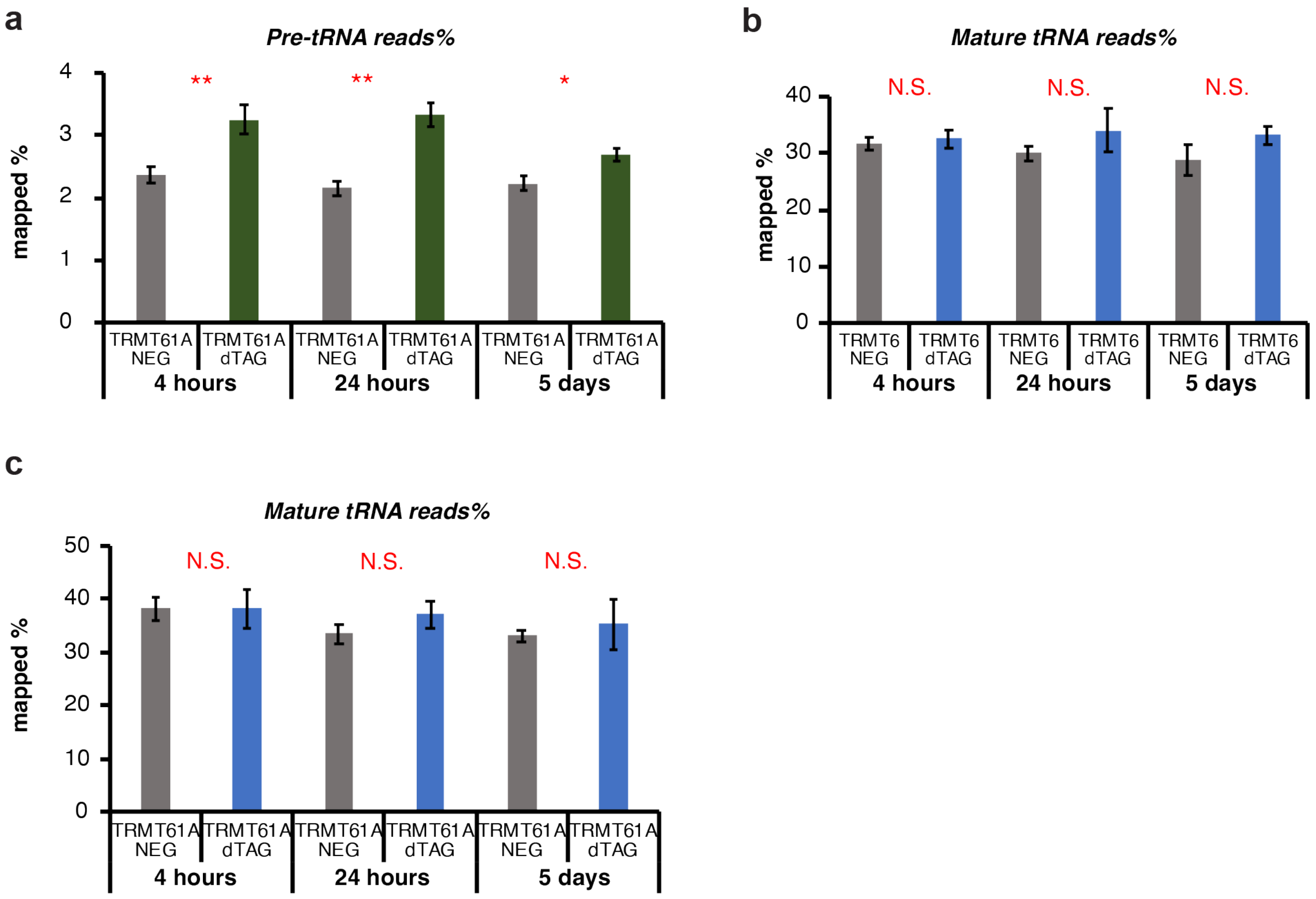

### Supplementary Figure 5.

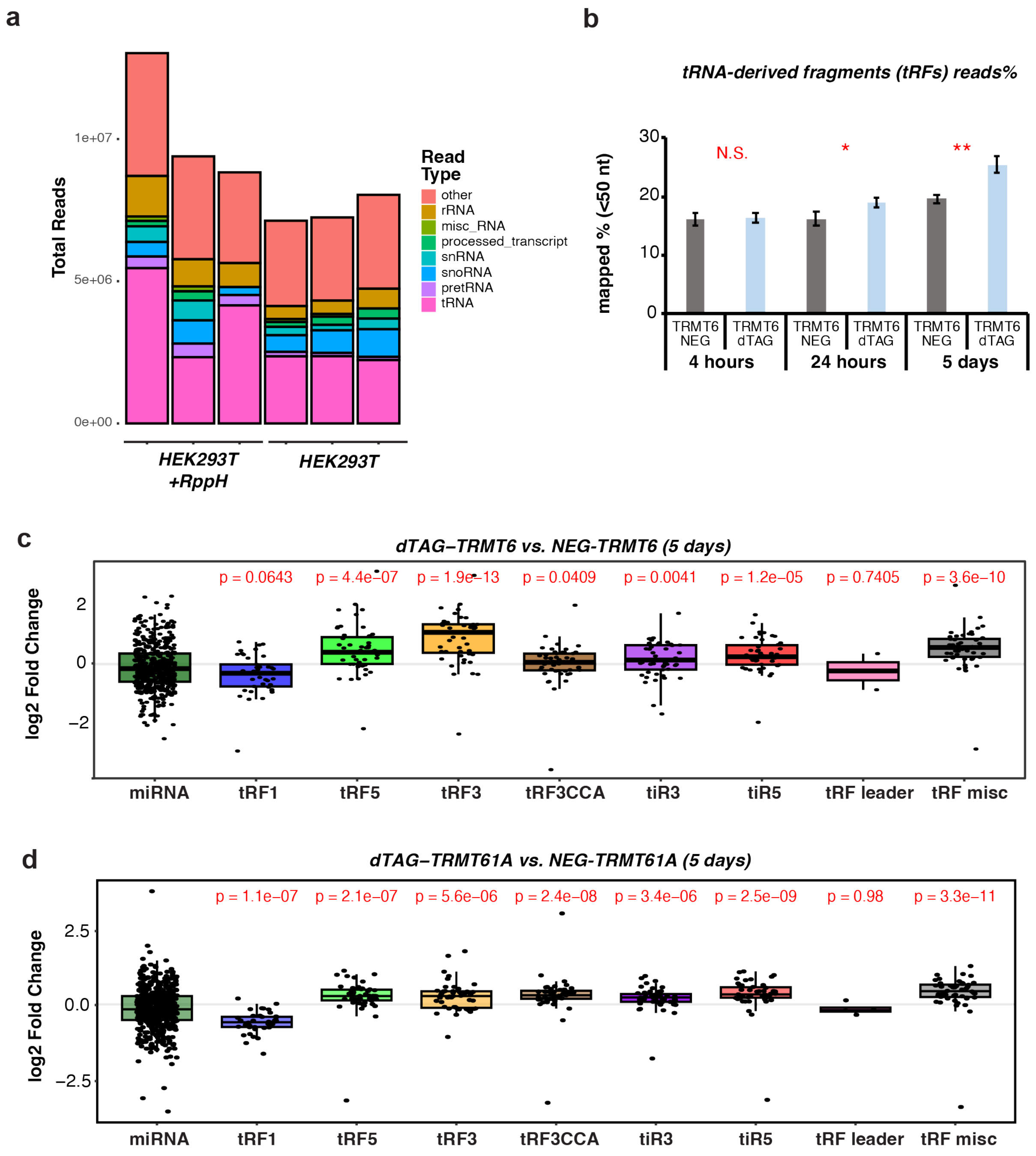

**Supplementary Figure 6.**

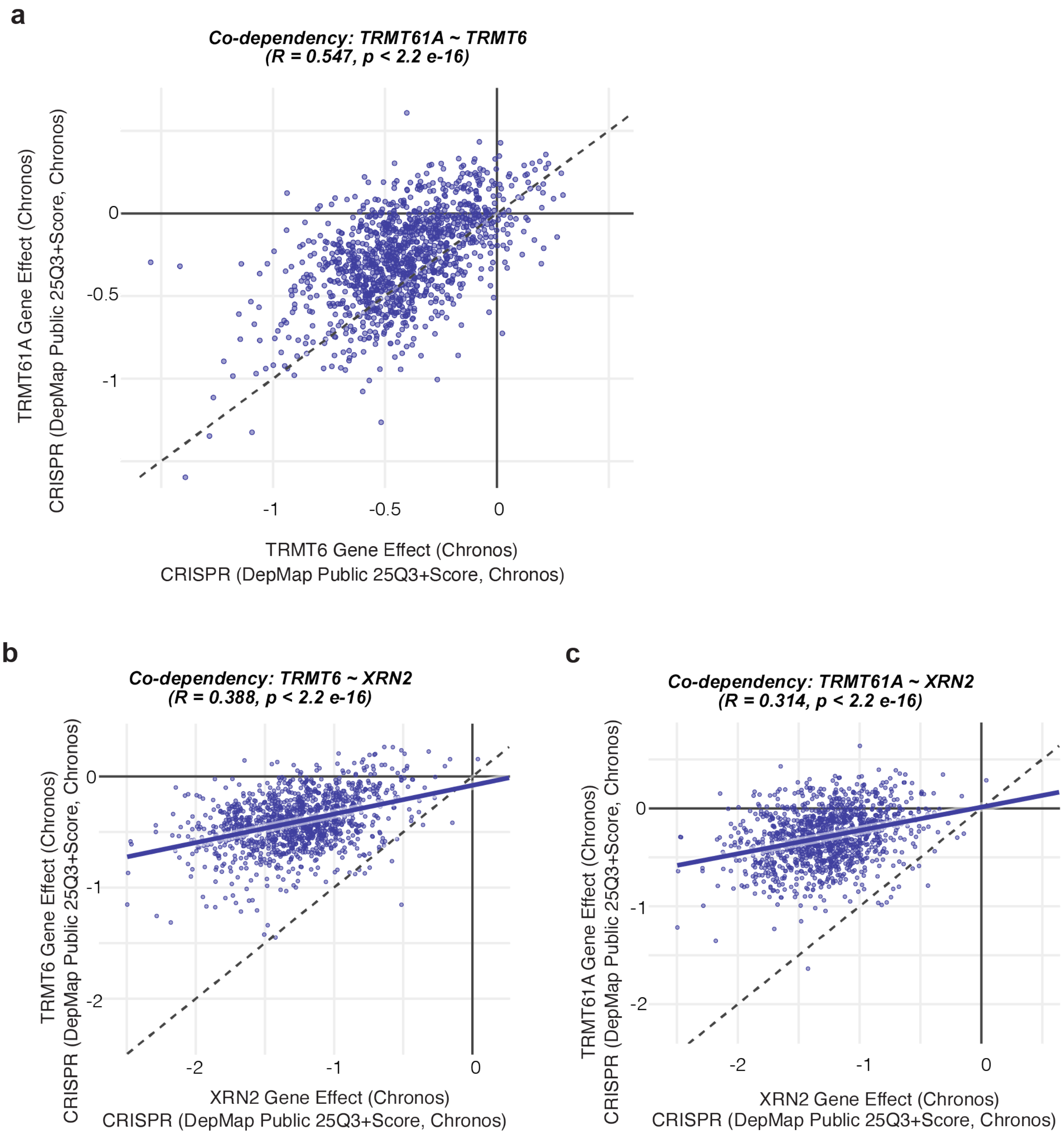
